# Computation in the human cerebral cortex uses less than 0.2 watts yet this great expense is optimal when considering communication costs

**DOI:** 10.1101/2020.04.23.057927

**Authors:** William B Levy, Victoria G. Calvert

**Author notes:** WBL conceptualized the study, developed the theoretical aspects and their description. WBL and VC developed the energy-audit and its description.

## Abstract

Darwinian evolution tends to produce energy-efficient outcomes. On the other hand, energy limits computation, be it neural and probabilistic or digital and logical. After establishing an energy-efficient viewpoint, we define computation and construct an energy-constrained, computational function that can be optimized. This function implies a specific distinction between ATP-consuming processes, especially computation *per se* vs action potentials and other costs of communication. As a result, the partitioning of ATP-consumption here differs from earlier work. A bits/J optimization of computation requires an energy audit of the human brain. Instead of using the oft-quoted 20 watts of glucose available to the brain (1, 2), the partitioning and audit reveals that cortical computation consumes 0.2 watts of ATP while long-distance communication costs are over 20-fold greater. The bits/joule computational optimization implies a transient information rate of more than 7 bits/sec/neuron.

**Significance Statement:** Engineers hold up the human brain as a low energy form of computation. However from the simplest physical viewpoint, a neuron’s computation cost is remarkably larger than the best possible bits/joule – off by a factor of 10^8^. Here we explicate, in the context of energy consumption, a definition of neural computation that is optimal given explicit constraints. The plausibility of this definition as Nature’s perspective is supported by an energy-audit of the human brain. The audit itself requires certain novel perspectives and calculations revealing that communication costs are 20-fold computational costs.

---

This paper examines neural computation from the perspective that Nature favors efficiency. To do so requires first quantifying a defined form of information generation that is common to evolved cortical computation. Second, we quantify cortical costs. Given the context of Darwinian fitness, a bits/joule optimization justifies our definition of computation as Nature’s perspective, as opposed to an *ad hoc* definition by engineers. Nevertheless, some effort is expended on aligning our definition with a particular first-principles derivation of energy-optimized computation arising from statistical mechanics.

Key functions in our investigation are ATP production and usage, processes which are dependent on glucose and oxygen. Thus, our energy optimized function can be expressed in terms of joules (J) per cycle, watts (W≡J/sec), moles of ATP, oxygen, or glucose per operation or per sec. Often, computer scientists, e.g. REFS, make a generic comparison between the power expenditure of computers vs the ≈20 watts of glucose consumed by the human brain. To further facilitate comparisons between the brain and engineered computers, we offer a partitioning of the human brain energy budget in a form homologous to traditional computing. The finding is that neural computation consumes 0.17 watts of ATP and cortical communication consumes 4.6 watts of ATP.

To measure computational costs requires a definition of computation. However taking the perspective of analog computation, any transformation qualifies as a computation. Likewise, any such transformation can be quantified using a variety of measures, arguably the most popular being Shannon’s mutual information (3). Without denying the acceptability of this most general perspective, we addend an additional property in our identification of neural computation: neural computation must be interpretable as some type of inference, e.g., the logical inference of digital computation or a Bayesian, statistical inference with evolution providing an implicit prior. In fact we identify commonalities between these two forms of computation: energy expenditures and the applicabilty of Shannon’s measurement.

Compared to the minimalist perspective of physics e.g.,(4), neural computation appears tremendously expensive. Because we assume Nature always gets it about right microscopically, the theory section concludes that the energy-optimized computation of physics is overly reductionist, and it must give way to a different, broader viewpoint. The suggested viewpoint is that Nature requires each neural system to deliver its computational information in a time-sensitive manner. To say it another way, a bits/J optimization of computation must heed a separate, minimal bits/sec requirement for regional brain computation. Thus, some scaled version of this bits/sec regional requirement will hold for individual neurons themselves. This additional constraint forces certain compromises on energy efficiency because energy-efficiency decreases as bits/sec increase, e.g. (5–7). As a result, the ultimate optimization must heed an unknown but requisite bits/sec. Rather than evaluate the organism’s information needs based on its fitness and niche, we take a bottom-up approach and consider how energy is used.

The approach uses empirical neuroscience to quantify various energy consuming processes. This forces us to combine various neuroanatomical, neurophysiological, and biophysical observations. With a well-defined form of computation, it is then possible to create a bits/J optimization even without knowing the required bits/sec. That is, because energy limits bits/sec of computation and communication, using the energy available for the bits/J optimization instructs the parameterization of the bits/sec calculation. Thus the bits/sec calculation is a prediction. The bridge between the bits/J optimization and the bits/sec calculation is *N* – the number of excitatory synaptic events required, on average, to reach threshold – and an important part of the Results proves this fact with precisely specified assumptions. One can also view the optimization result as a consistency check on the energy-audit since an empirically based, inferred value for *N* is part of the audit’s development. Before developing these results, we consider the best possible bits/J that physics offers and map this result into the context of a simple neural computation.

## Results

### Relating neural computing to the maximum efficiency of irreversible computing

Our approach to quantifying and interpreting the energetic cost of neural computation, and its optimization, is inspired by the physical limits on irreversible computation. Still we do not stray too far from what theoretical neuroscience has had to say about measuring information, now in the context of computation. For the sake of our initial comparison, suppose a neuron’s computation is just its transformation of inputs to outputs. Then, quantifying the information passed through this transformation (bits per sec) and dividing this information rate by the power (W = joules/sec) needed for such a transformation yields bits/J. This ratio will be our efficiency measure. In neuroscience, it is generally agreed that Shannon’s mutual information (*MI*) can be used to measure something about the bit-rate of neural information processing, neural transformations, or neural communication, e.g., (8–15). Specifically, we use mutual information and an associated rate of postsynaptic energy-use, which will allow a comparison with the optimal bits/J for computation as developed through physical principles. To understand this analogy with the result of statistical mechanics, assume the only noise is wideband thermal noise, k𝒯 ≈ 4.3 · 10^−21^ J (Boltzmann’s constant times absolute temperature, 𝒯 = 310 K). The bits/J ratio can be optimized to find the best possible energetic cost of information, which is (*k𝒯* ln 2)^−1^.

To give this derivation a neural-like flavor, suppose a perfect integrator with the total synaptic input building up on the neuron’s capacitance. Every so often the neuron signals this voltage and resets to its resting potential. Call the signal *V*_*sig*_, and rather unlike a neuron, let it have mean value (resting potential) of zero. That is, let it be normally distributed 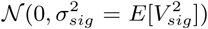. The thermal noise voltage-fluctuation is also a zero-centered normal distribution, 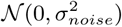. Expressing this noise as energy on the membrane capacitance, 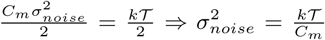 (16–18). Then using Shannon’s result, e.g., theorem 10.1. 1 as in (19), the nats per transmission are 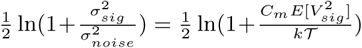 (with natural logarithms being used since we are performing a maximization). Converting this to bits, and calling this result the mutual information channel capacity, 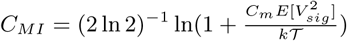.

Next we need the energy cost, the average signal joules transmission developed on the fixed *C*_*m*_ by the synaptic activation, 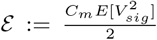. Dividing the bits/sec *C*_*MI*_ by the J/sec *ε* yields the bits-per-joule form of interest; 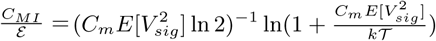. This ratio is recognized as the monotonically decreasing function 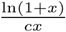 with *x, c* > 0. Therefore maximizing over 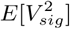 but with the restriction 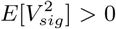 is a limit to the left result, an approach to zero bits/sec. That is,

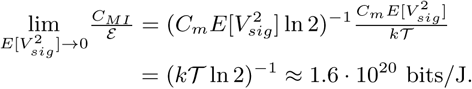

Two comments seem germane. First, physicists arrived at this value decades ago in their vanquishing of Maxwell’s demon and its unsettling ability to create usable energy from randomness (4). In their problem, the device (the demon) is not obviously computational in the neural sense; the demon just repeatedly (i) senses, (ii) stores, and (iii) operates a door based on the stored information, and then (iv) erases its stored information as it continues to separate fast molecules from the slower ones (20, 21): see Fig 1. Moreover, even after simplifying this cycle to steps (i), (ii) and (iv), physicists do see the demon’s relevance to digital computation. Such a cycle is at the heart of modern computers where computation occurs through repetitive uses, or pairwise uses, of the read/write/erase cycles. For example, bit-shifting as it underlies multiplication and the pairwise sensing and bit-setting (then resetting) of binary, Boolean logical operations reflect such cycles. Thus, as is well known from other arguments e.g., (4), (22), the limit-result of physics sets the energy-constraining bound on non-reversible digital computation. Regarding (iii) it would seem that if the demon communicates and controls the door as slowly as possible (i.e, the limit of time going to infinity), there is no need to assign an energy-cost to these functions.

**Fig. 1.**
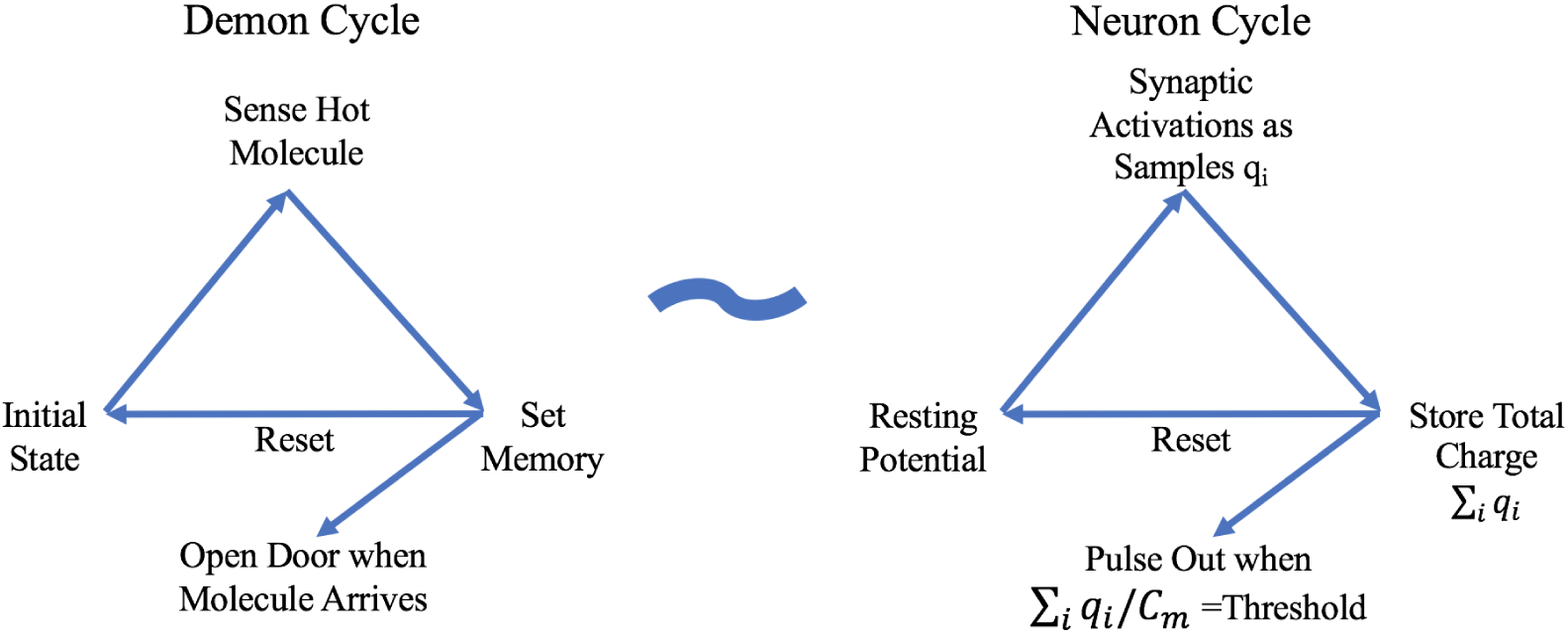
Maxwell’s demon cycle is analogous to the neuron’s computational cycle. The initial state in the demon cycle is equivalent to the neuron at rest. The demon sensing a fast molecules is analogous to the synaptic activations received by the neuron. Whereas the demon uses energy to set the memory and then opens the door for the molecule, the neuron stores charge on the membrane capacitance (C_m_) and then pulses out once this voltage reaches threshold. Simultaneous with such outputs, both cycles then reset to their initial states and begin again. Both cycles involve energy being stored and then released into the environment. The act of the demon opening the door is ignored as an energy cost; likewise, the neuron’s computation does not include the cost of communication. Each q_*i*_ is a sample and represents the charge accumulated on the plasma membrane when synapse *i* is activated.

Secondly, compared to the estimates here of a neuron cycling from reset to firing to reset, this physics result is unimaginably more efficient. Suppose that the computational portion of a human cortical neuron has capacitance *C*_*m*_ ≈ 750 pF (obtained by assuming the human neuron’s surface area is a about three times a rat’s pyramidal value of 260 pF (23)) and suppose this neuron resets to *V*_*rst*_ = −0.066 V while firing threshold is *V*_*θ*_ = −0.050 V. Then in the absence of inhibition, the excitatory synaptic energy needed to bring a neuron from reset to threshold is 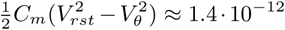 J/spike. Assuming 4 bits/spike, the bits/joule are 2.9 · 10^12^. Compared to the optimal limit set by physics, this efficiency value is 10^8^ times less energy-efficient, a seemingly horrendous energy-efficiency for a supposedly optimized system.

#### The disagreement reorients our thinking

In the context of understanding neural computation via optimized energy-use, this huge discrepancy might discourage any further comparison with thermal physics or the use of mutual information. It could even discourage the assumption that Nature microscopically optimizes bits/J. But let us not give up so quickly. Note that the analogy between the four-step demon versus an abstract description of neural computation for one interpulse interval (IPI) is reasonable (see Fig 1). That is, (i) excitatory synaptic events are the analog of sensing, these successive events are (ii) stored as charge on the plasma membrane capacitance until threshold is reached, at which point (iii) a pulse-out occurs, and then (iv) the “memory” on this capacitor is reset and the cycle begins anew. Nevertheless, the analogy has its weak spots.

The disharmony between the physical and biological perspectives arises from the physical simplifications that time is irrelevant and that step (iii) is cost-free. While the physical simplifications ignore costs associated with step (iii), biology must pay for communication at this stage. That is, physics only looks at each computational element as a solitary individual, performing but a single operation. There is no consideration that each neuron participates in a large network or even that a logical gate must communicate its inference in a digital computer in a timely manner. Unlike idealized physics, Nature cannot afford to ignore the energy requirements arising from communication and time constraints that are fundamental network considerations (24) and fundamental to survival itself (especially time).

According to the energy audit, the costs of communication between neurons outweighs computational costs. Moreover, this relatively large communication expense further motivates the assumption of energy-efficient IPI-codes (i.e., making a large cost as small as possible is a sensible evolutionary prioritization). Thus the output variable of computation is assumed to be the IPI, or equivalently, the spike generation that is the time-mark of the IPIs endpoint.

Furthermore, any large energy cost of communication sensibly constrains the energy allocated to computation. Recalling our optimal limit to the left (i.e., the asymptotic zero bits/sec to achieve the (*k𝒯* ln 2)^−1^ bits/J), it would be unsustainable for a neuron to communicate minuscule fractions of a bit with each pulse out. To communicate the maximal bits/spike at low bits/sec leads to extreme communication costs because every halving of bits/sec requires at least a doubling of the number of neurons. Such an increasing number of neurons eventually requiring longer and wider axons; thus intuition says using more neurons at smaller bit-rate leads to a space problem. Such a space problem is generally recognized as severely constraining brain evolution and development as well as impacting energy-use (25–30). It is better for overall energy consumption and efficiency to compute at some larger, computationally inefficient bits/IPI that will feed the axons at some requisite bits/sec, keeping neuron number and neuron density at some optimal level. To say it another way, a myopic bits/J optimization can lead to a nonsense result, such as zero bits/sec.

*Nevertheless, assuming efficient communication rates that go hand-in-hand with the observed communication costs, there is still reason to expect that neuronal computation is as energy-efficient as possible in supplying the required bits/sec of information to the axons*. The problem then is to identify such a computation together with its bits/J and bits/sec dependence.

### Information rate estimates and optimizing computation under energy constraints

We assume a neuron is constructed to estimate the value of a particular scalar latent random variable based on the rate of its net synaptic excitation. The computation is an implicit probabilistic inference via Lindley-Shannon-Bayes (see below and (31)). Specifically, the IPI, a sufficient statistic, is the time it takes for net synaptic excitation to move the membrane potential from reset to threshold. In other words, a neuron’s computation is the process where it adds together synaptic inputs over time, and this time is implicitly an estimate of the value of the neuron’s latent variable.

#### The components and the assumptions

This section derives an energy-optimization result for neuronal computation that acknowledges a specific subset of energy-uses. By incorporating all the appropriate energy costs into the bits/J function, there is an implicit enforcement of a positive bits/sec. That is, the limit result implying zero bits/sec is avoided. To reach this new optimization requires the introduction of some definitions, a few assumptions, and then the function that quantifies energy efficiency (31).

Assume a neuron is built to estimate, in an unbiased fashion, a scalar random variable (RV) Λ. This estimate is the RV 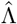 and is based upon the intensity of the neuron’s excitatory input activations. Although a neuron never calculates its information rate, we must. Thus, assume Nature has an implicit prior distribution, and assume that this prior is a probability density, *p*(*λ*), that is continuous and possess a finite mean, *E*[Λ]. By virtue of energy-efficiency, IPI coding is used; such coding implies that the output of a neuron is a time-interval, the RV *T*. That is, an IPI code is assumed because of its high energy-efficiency (we know of none better that uses *{*0, 1*}* coding) and because constant amplitude pulses imply that all information is in the IPIs. Assume the excitatory synaptic activations consume energy in a linear fashion. Thus the average synaptic energy consumption is proportional to *E*[Λ · *T*], the expected value of the product of the random input intensity to the neuron times the random duration of the first IPI. We are now most of the way toward specifying the sense in which the energy devoted to computation is optimal.

Assume that the uncertainty of a neuron’s estimation, as coded by the time of action potential (AP) generation at the initial segment, far exceeds any uncertainty caused by axonal jitter. Then a bits/J function (much like in (5)) will be constructed. However, instead of optimizing the axon’s spike-rate as a function of the components of axonal energy-use, here the number (*N*) of synaptic excitations per IPI is optimized as a function of specified energy consumers including the energy-use associated with APs and the events they trigger. The function to be maximized is 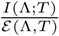 where the numerator is the information generated by the neuron’s computation for its first IPI with no axonal information loss. The energy-use per IPI, ε (Λ, *T*) takes into account the energy devoted to (i) communication, (ii) computation, and (iii) *Other*, which encompasses the combined energy for AP-triggered maintenance and synaptic modifications (this last includes, *inter alia*, receptor-modification, metabotropic synaptic activations, synaptogenesis, and all the cell biology needed to support these processes). This ratio of expectations is concave in synaptic activations per IPI, and thus the bits/J can be maximized. As the corollaries of the next subsection make clear, the energy devoted to computation restricts the precision of a neuron’s estimation and restricts the information a neuron generates.

The value of *N* that optimizes 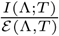 is a consistency check for the energy-audit values. In particular, this optimization might agree or disagree with the 2500 estimate (a function of the number of input lines, synaptic failure rates, and the assumption that the average firing rate of each input to a neuron equals the average firing rate of the neuron).

### Valuing Lindley’s information gain for the first IPI

Using maximum entropy and its ability to produce optimal probability distributions (e.g., minimax mean squared error (MSE), (32)), (31) infers an optimal form of the likelihood, *p*(*t* | *λ*). Here *t* is the IPI whose inverse is directly proportional to 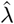. Also, (31) proves the sufficiency of a specific form of the marginal *p*(*λ*) while conjecturing necessity. These results hinge on the constraints of energy-use and unbiased estimation. Upon inspection, the resulting distributions can be used in Lindley’s information-gain formulation. That is, (33) demonstrates that Shannon’s mutual information measures the information gain of a Bayesian who uses experimental measurements to update his prior to a posterior distribution.

From equations 12 and 6 of (31), the optimization results are 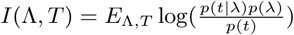, with 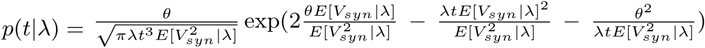. While the only consistent marginal distribution we have yet to discover is, 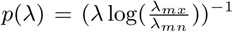 with 0 *< λ*_*mn*_ *< λ < λ*_*mx*_ *< ∞*.

#### Random variation of synaptic activations dominate the estimation error

It is worth pausing at this point to note that the variance is directly proportional to the mean rate of arrival of synaptic excitations. This variance 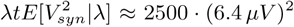 is a denominator term in the exponential part of *p*(*t* | *λ*), and it arises from the signal itself. When *N* is 2500, this signal variance is the dominant randomization, overshadowing other forms of noise. Specifically using the earlier thermal noiselevel result, 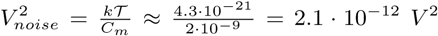, (or as a neurophysiologist might better appreciate, the standard deviation of this noise is 1.45 *µ*V). The squared value, as energy on the membrane capacitor, is small compared to the energy needed to reach threshold.

Shot-noise is small but might reasonably be included in an information calculation (34). As developed in Methods and based on biophysical simulations (23), the initial-segment NaV 1.6 shot-noise is less than 10% of the synaptic randomization, ≈250 events under slow depolarization where the event amplitudes are about the same size as 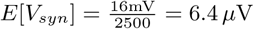. To account for this initial segment noise around threshold, one can increase the variance term of the drifted diffusion (see Methods). However, the effect is small (0.12 bits) and is ignored in what follows. In sum, the dominant source of information degradation for IPI-coding is the imprecise clocking of neural networks and the random arrival of synaptic excitations to a neuron.

#### Simplifying the distributional forms, calculating error, and determining information rates

Closing in on the bits/J optimization, we simplify the conditional probability density. Assume an empirical distribution of synaptic weights such that the second non-central moment is equal to twice the mean squared (e.g., an exponential distribution). Note also that *θ* can be written as the product *N*, the average number of synaptic increments, multiplied by the average synaptic incrementing event *E*[*V*_*syn*_ | *λ*] (with inhibition and capacitance taken into account, (31)). That is, *θ* = *N E*[*V*_*syn*_ | *λ*]. Putting this assumption to work, a simplification obtains, and there are two new corollaries related to the above optimal probability distributions.

##### Lemma 1

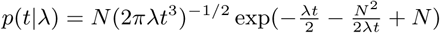.

*Proof*: Start with *p*(*t*|*λ*) given earlier, substitute using *θ* = *N E*[*V*_*syn*_ | *λ*], and then note that 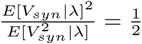.

At this point there is an instructive and eventually simplifying transform from *p*(*t* |*λ*) to the distribution of the estimate that the neuron is implicitly creating, 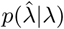. The transform is defined by the unbiased requirement that is one of the constraints producing the earlier optimization results (31). Given that the relationship between *θ* and *N* is now 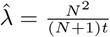 or equivalently 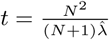,

##### Lemma 2a

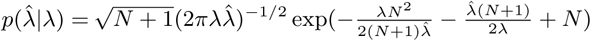.

##### Lemma 2b

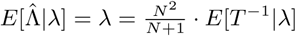;

##### Corollary 1

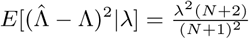.

*Proofs*. See Methods.

As Lemma 2b shows, the estimate is indeed unbiased, and as the corollary shows, devoting more energy to computation by increasing *N* reduces the error of the estimation. Equivalently, as *N* grows, the standard deviation decreases at the rate of 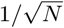. Of course, computational costs increase in direct proportion to *N*.

This corollary adds additional perspective to our definition of a neuron’s computation as an estimation. Furthermore, the new likelihood, 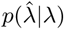, is particularly convenient for calculating information rates, a calculation which requires one more result. That result is the marginal distribution of 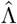. Because the only known sufficient density (and arguably the simplest) is 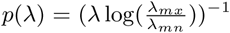, the estimate’s marginal density is simply approximated via

##### Lemma 3

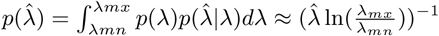, where the approximation arises by the near identity of the integral to *p*(*λ*) assuming the range of *λ* and 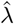 is the same. Moreover, the lack of 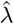 bias for all conditioning values of *λ* hints that the approximation should be good. In fact, a naïve numerical evaluation of 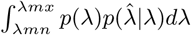 indicates zero difference between this integral and 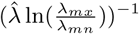; however, see SI for a more precise analysis of this approximation.

The information rate per first-IPI can now be evaluated.

##### Corollary 2

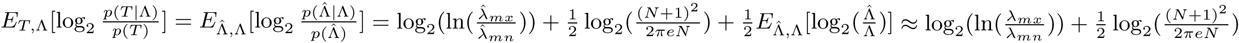

*Proof*: 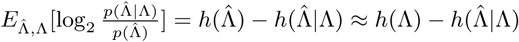.

This is a good approximation because of the near equivalence of any marginal expectations of the two marginal densities compared above. This approximation produces a value of ca. 6.94 bits when *N* = 2500 and 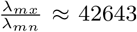, a ratio consistent with a spontaneous synaptic transmission of 1 Hz over 10^4^ synapses and with an average firing rate of 1.6 Hz (see Methods ans SI). Therefore by this result, the bit-rate increases with the number of synaptic activations per IPI essentially at the anticipated rate of 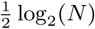. However, this nearly seven bits per IPI is an upper bound, and at this stage of the development of Lindley’s measure, we downgrade the bits/IPI estimate to 4.6 bits/IPI (see Methods for details), i.e., almost 7.4 bits/sec. The principle cause of this downgrading is the small bit rate of IPIs succeeding the first IPI and the naive, fixed threshold model currently being used.

#### The bits/J optimization confirms the assumed value of N in the energy-audit

Finally there is enough to perform an optimization of the computational bits/J. Doing so asks if the values and assumptions of the energy audit are consistent. In particular, the following confirms that the above use of *N* = 2500 is very close to an appropriate value. Taking *N* as an optimizable variable and dividing the information rate per IPI by energy-use per IPI yields the following ratio with units of bits/J for one neuron: 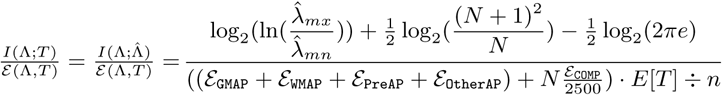.

with *E*[*T*] *÷ n* = 4 · 10^11^ rescaling the cortical energy to one neuron and one IPI.

The energy-consumption function is notably different from the usual form as it specifically does not include energy consumption that grows linearly with time, e.g., axonal resting potential costs (see (31) where such costs are assumed to be borne by the system decision-maker). Because its denominator is scaled to J/IPI/neuron, this energy function includes the cost of just the one output AP at the end of the IPI. The AP cost, as estimated in the energy audit, consists of four components, the GM axons, the WM axons, the relevant presynaptic functions, and a fraction of *Other*; i.e., resp. (ℰ_GMAP_+ ℰ_WMAP_+ ℰ_PreAP_+ ℰ_OtherAP_) = (0.75+0.54+0.19+0.07) = 1.54 W. The derivation of these values is detailed in the next section and Methods, including *ℰ*_COMP_ = 0.17 W..

For these joule-costs, the bits/J optimization calculation yields N ≈ 2428, missing the assumed value of 2500 by 3%. In fact, increasing AP-costs by ca. 3% or decreasing computational costs by ca. 3% yields the desired value of the 2500. Even we must believe such nearly perfect agreement is, to some extent, fortuitous. To be clear, the energy-audit was performed before this optimization calculation, and there was no recursive tuning of the audit to get such a close agreement between values. In fact, we have been using 2500 for *N* for quite some time, well before doing this optimization; moreover, 2500 can arise many ways, e.g., 12,500 synapses per neuron and a failure rate of 80%. Fig 2 illustrates the differing sensitivities of this result to the two distinct energy dependencies of the optimization; Fig 2A is computational energy vs Fig 2B, which is AP-associated non-computational energy. Moreover, the consistency result has some flexibility. Increasing or decreasing AP costs while decreasing or increasing computational costs by the same percentage, respectively, leaves the inferred *N* value unchanged. Finally, this optimization does not incorporate the ratio of the constant, time-proportional costs which include the largest ATP-consumer, axonal resting potentials.

**Fig. 2.**
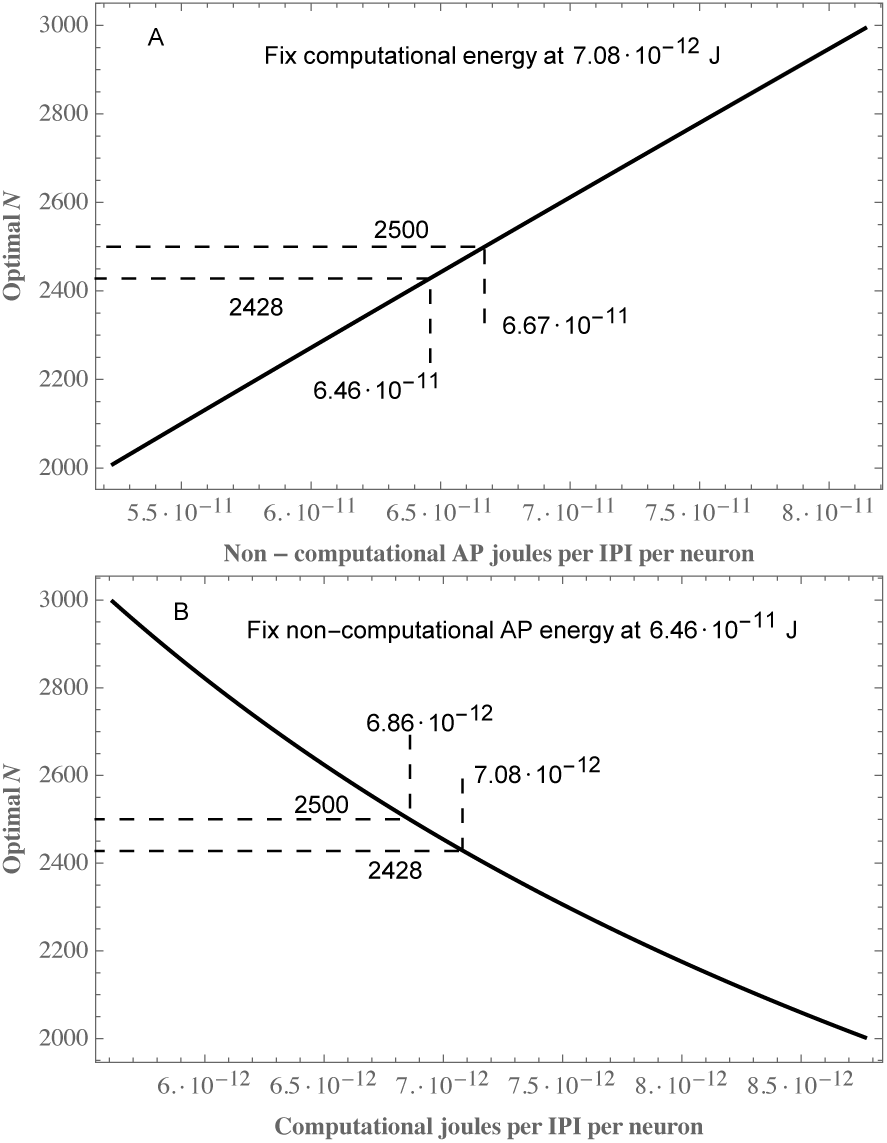
Near exact consistency between the energy-audit’s *N* = 2500 and the optimization implied value *N* = 2428. The plotted curves indicate the sensitivity of the optimization result to energy allocations. Perfect consistency (*N* = 2500) requires either (A) increasing non-computational AP energy-use from 6.46 to 6.67 · 10^−11^ J/neuron/IPI, (B) decreasing the computational energy budget from 7.08 to 6.86 · 10^−12^ J/neuron/IPI, or (C) some even smaller alterations of both energy consumers. Because *N* is so large, the curvature in A is imperceptible.

### Energy audit

#### ATP use for computation and communication

The values of this section preceded the calculations of the previous section. Here, data from empirical neuroscience are employed to estimate the joules per second devoted to the microscopic processes used by the previous section. To accomplish this goal, divide cortical processes into computational costs (i.e., postsynaptic costs) and communication costs (axonal and presynaptic costs). Since ATP is the molecule used by energy-consuming processes in brain, the energy consumption of each process is based on ATP consumption as in (35), thus the term ATP-watts.

As derived below, computation consumes less than 0.2 ATP-watts or less than *one one-hundredth of the nominal and oft quoted 20 watts that would be produced by complete oxidation of the glucose taken up by the brain* (1). Fig 3 compares cortical communication costs to computational costs. For some, the rather large cost of communication might be surprising. The lion’s share of these ATP-watts goes to communication where it contributes to signal velocity and information rates (36, 37). Combining gray matter (GM) communication costs with the total white matter (WM) costs accounts for 93% of the total ATP-watts compared to 3.4% for computation. Supposing, rather generously, that WM *Other* consumes 0.65 W, then GM plus WM communication accounts for 87% of the ATP-W, thus giving a ratio of 25:1 for communication vs computation. Because so little energy goes to computation and because so much goes to the axonal resting potential, the ratio just calculated is particularly sensitive to average firingrate. Specifically, computational costs increase much more quickly with firing rate than the total cost of communication.

In what follows the reader will find progressively more details explaining the values in Table 1 and Fig 3, including the sensitivity of computational costs to firing rate, the derivation of computational costs, and finally the derivation of communication costs. Even more details can be found in the Methods section. These details include the underlying calculations and accompanying assumptions.

**Table 1.** Rudimentary partitioning, glucose to ATP See Methods and SI Tables for details and citations

| Brain/Region (weight) | Watts (complete oxidation) | Unoxidized (equivalent watts) | Heat watts | ATP-watts |
| --- | --- | --- | --- | --- |
| whole brain (1495 g) | 17.0 | 1.86 | 8.89 | 6.19 |
| cerebellum (154 g) | 1.77 | 0.19 | 0.93 | 0.65 |
| other regions (118 g) | 1.65 | 0.18 | 0.87 | 0.60 |
| forebrain cortex (1223 g): |  |  |  | 4.94 |
| white (590 g) | 5.07 | 0.56 | 2.66 | 1.85 |
| gray (633 g) | 8.45 | 0.93 | 4.43 | 3.09 |

**Fig. 3.**
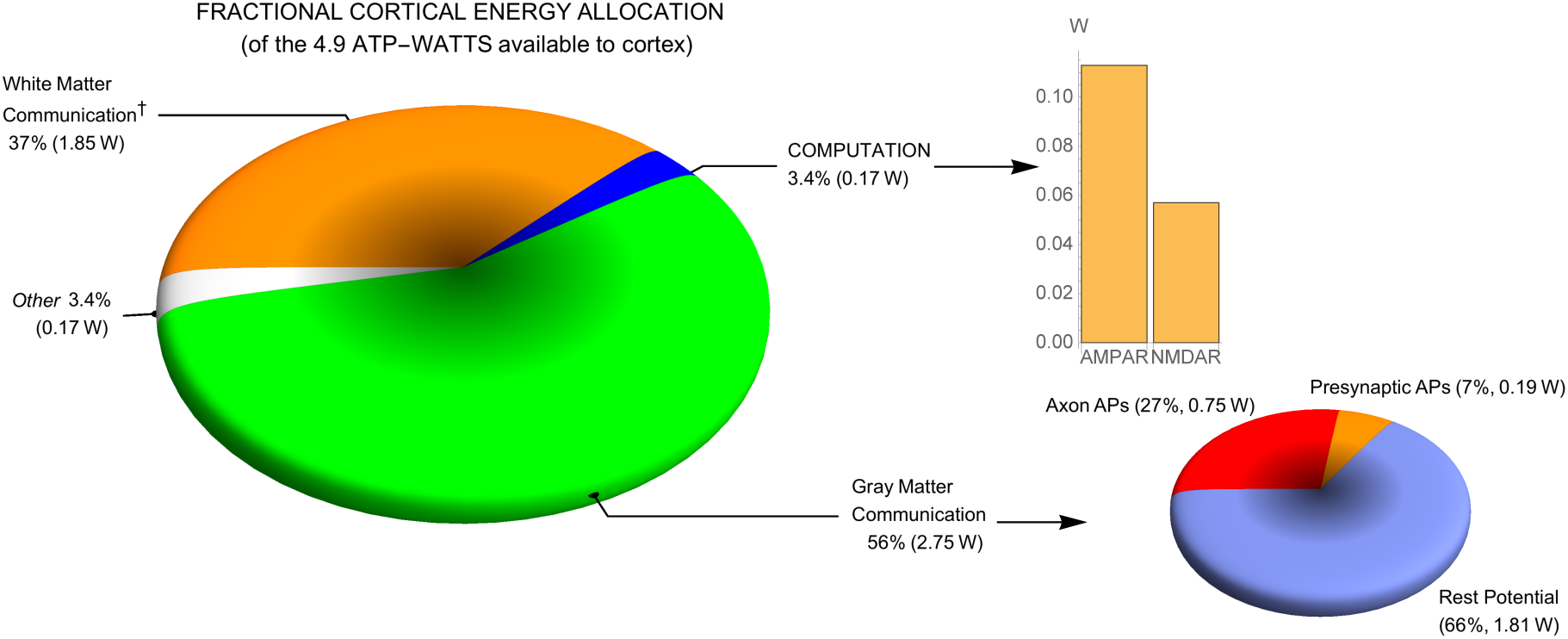
Computation costs little compared to communication. GM communication alone accounts for more than half of the cortical energy use (big pie chart). Computation, the smallest consumer, is subpartitioned into the two ionotropic glutamate receptors (bargraph). *Other* includes synaptic modification and maintenance. The small pie chart sub-paritions GM communication after re-scaling (2.75 W =100%). See Results, Table 1, and Methods for details. ^*†*^WM communication includes its *Other* in addition to resting and action potentials.

#### An energy-use partitioning based on glucose oxidation

Two approaches are used to evaluate the energy consumed by the brain: a top-down partitioning of glucose-watts converted to ATP-W and a bottom-up series of biophysical calculations based on ATP-use by partitioned aspects of a functioning neuron. Table 1 and Fig 3 summarize the results of both approaches. The 17 W of glucose potential energy from recent PET scan research (see Table S1 (38)) replaces Sokoloff’s 20+ W from the 1950s. The PET scan research presents regional per-gm values, and these values are scaled by the brain mass partitioning of (39). Of this total glucose uptake, ca. 11% is not oxidized (40) although quantitative conclusions from scanning studies are challenged by arteriovenous blood differences that obtain a smaller non-oxidized fraction (see Supplement). After removing the 8.89 W that go to heating, there are only 6.19 ATP-W available to the whole brain. Regional partitioning whittles this down to 3.09 ATP-W for the categories of computation, communication, and *Other* of the cerebral gray matter. By *a posteriori* design, the 1.6 Hz average firing rate is chosen to match the 3.09 available ATP-watts when *Other* (synaptic modification and maintenance) is valued the same as computational costs. When 2.5 Hz is used and *Other* energy-use is again matched to computational costs, the total of 3.79 W exceeds the 3.09 by ca. 17% of this nominally available value.

#### Firing Rate

In regard to average firing rate, one might initially guess a value of one pulse per neuron per decision-making interval (DMI); that is, one pulse per visual fixation, which is the time it takes to decide where to aim the next saccade. This would be about one pulse per 400 msec in humans, implying 2.5 Hz for an average firing rate.

Although (41) prefers a human average firing rate closer to 0.1 Hz than 1 Hz, our preferred estimate is 1.6 Hz, arising from the book-balancing argument just above. Then using our bottom up calculation for the excitatory postsynaptic ion-flux per AP per neuron, 1.6 Hz combined with the number of neurons exactly accounts for the 0.17 W available. The linear relationship between firing rate and energy consumption (Fig 4) has a substantial baseline energy consumption of 1.81 W (y-axis intercept). This intercept includes 0.01 W of *Other*. More important is the 1.8 W arising from resting axon conductance required for resting potential and stable behavior (42). In the case of the dendrite, computational costs are zero at zero firing rate, a theoretical limit result which, as argued earlier, is a nonsense practical situation. Dendritic leak is assumed to be essentially zero since we assume, perhaps controversially (cf. (35)), that a cortical neuron is under constant synaptic bombardment and that all dendrosomatic conductances are due to synaptic activation and voltage-activated channels. That is, a neuron resets after it fires and immediately starts depolarizing until hitting threshold.

**Fig. 4.**
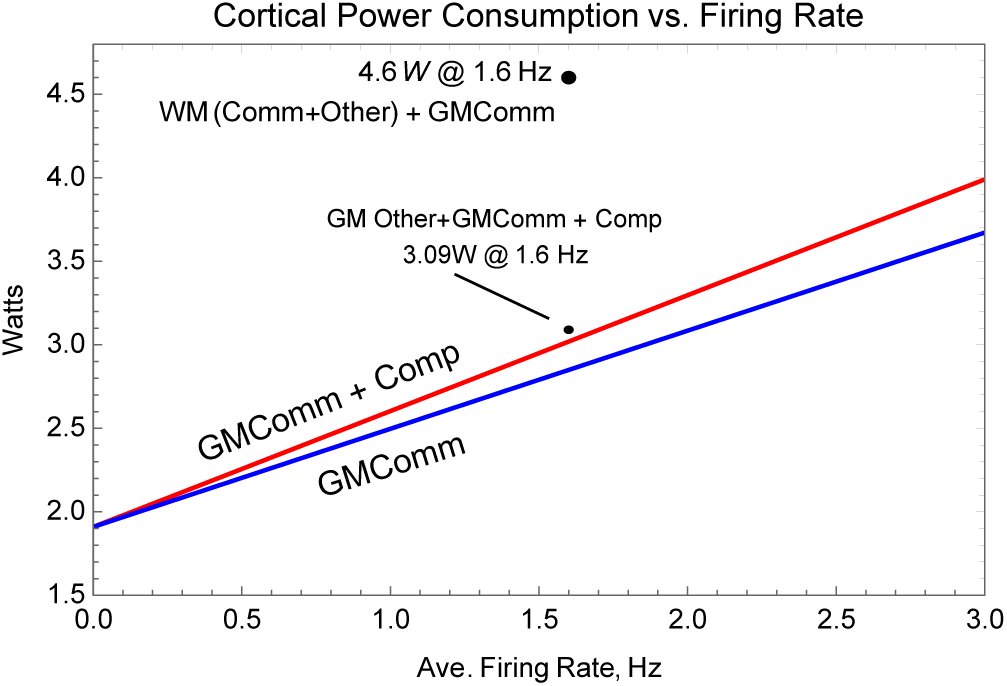
Energy-use increases linearly with average firing-rate, but for reasonable rates, computation (Comp) costs much less than communication (Comm). Comparing the bottom (blue) curve (GM communication costs) to the top (red) curve (GM communication cost plus computational costs), illustrates how little computational costs increase relative to communication costs. The large y-intercept value is 1.8 W for resting potential plus 0.1 W for a constant consumption by *Other*. The small point labeled GMAPOther+GMComm+Comp adds 0.07 W of AP-dependent *Other* to the GMComm+Comp curve, 0.17 W + 2.75 W @ 1.6 Hz. The large point, labeled WM(Comm + Other) + GMComm, shows the value of the combined communication cost, i.e., cortical GM at 1.6 Hz plus the total cortical white matter (WM) cost. See Table 1, and Methods for further details.

Computational costs are very sensitive to failure rates, which for this figure are fixed at 75%, whereas communication is only slightly sensitive to the synaptic failure rate (see below for more details).

#### Computation costs in the human brain

The energy needed to recover ion-gradients from the total excitatory synaptic current-flows/IPI determines the cost of computation for that IPI. Various quantitative assumptions feeding into subsequent calculations are required (see Methods and Supplement), but none are more important than the generic assumption that the average firing-rate of each input to a neuron is the same as the average firing-rate out of that neuron. Via this assumption, and assuming 10^4^ synapses per neuron and a 75% failure rate, the aggregate effects of inhibition, capacitance, and postsynaptic K^+^ conductances are implicitly taken into account. This aggregation is possible since increases of any of these parameters merely lead to smaller depolarizations per synaptic activation but cause little change in synaptic current flow per excitatory synaptic event. Indeed, such attenuating effects are needed to make sense of several other variables. A quick calculation helps illustrate this claim.

After taking quantal synaptic failures into account, substantial inhibition is required if there are to be 2500 excitatory events propelling the 16 mV journey from reset to threshold. That is, with 10^4^ inputs and the 75% failure rate, 2500 synapses are activated per IPI, on average. Activation of AMPARs and NMDARs provides an influx of three Na^+^’s for every two K^+^ that flow out. With an average total AMPAR conductance of 200 pS, there are 114.5 pS of Na^+^ per synaptic activation (SA). Multiplying this conductance by the 110 mV driving force on Na^+^ and by the 1.2 msec SA duration yields 15.1 fC per SA. Dividing this total Na^+^ influx by 3 compensates for the 2 K^+^ that flow out for every 3 Na^+^ that enter; thus, the net charge influx is 5.04 fC/SA. We assume that the voltage-activated, glutamate-primed NMDARs increases this net flux by a factor of 1.5, yielding 7.56 fC/SA (see Methods and SI Tables 3, 4, and 5 for more details and the ATP costs). Taking into account the 2500 synaptic activations per IPI yields 18.9 pC/IPI. Using a 750 pF value for a neuron’s capacitance, this amount of charge would depolarize the membrane potential 25.2 mV rather than the desired 16 mV. Clearly, the excitatory charge influx must be opposed by inhibition and K^+^ conductances to offset the total 7.56 fC net positive influx. Most simply, just assume a divisive inhibitory factor of 1.5. Then the numbers are all consistent, and the average depolarization is 6.4 *µ*V per synaptic activation. Because each net, accumulated charge requires one ATP to return the three Na^+^’s and 2 K^+^’; thus, the computational cost of this 16 mV depolarization is 7.1 · 10^−12^ J/neuron/spike, upholding the earlier approximation. In other words, the computational power required per cortex per spike is 0.17 W using 1.5 · 10^10^ neurons firing at a rate of 1.6 Hz. See Discussion and Methods for more remarks and explications.

#### Communication costs

As quantified in Methods and summarized in SI Tables 3 and 5, the GM long-distance communication cost of 2.75 W includes the partitioned costs of axonal resting potential, APs, and presynaptic transmission (neurotransmitter recycling and packaging, vesicle recycling, and calcium extrusion). The neurotransmission costs assume a 1.6 Hz firing rate and a 75% failure rate. From there, we use the (43) calculation that assumes one vesicle is released per non-failed AP. Differing from (43) while closer to earlier work (35), we assume that there is the same Ca-influx with every AP (44). Furthermore, we also use a more recent measurement of Na^+^-K^+^ overlapping current flows of the axonal AP (45). Of all the difficult but influential estimates here, none is more challenging and important than axonal surface area. See Methods for more details.

## Discussion

The Results contribute to our understanding of computation in the brain from the perspective of Nature. Essentially, the Results present a defined form of neural computation that is based on postsynaptic activation and that is (ii) a probabilistic inference. From this defined perspective, the corresponding optimal bits/J is calculated. In a meaningful sense, this calculation confirms the consistency of our numbers. The optimizing *N*, the average number of synaptic excitations underlying a neuron firing, is the value of *N* that results from our biophysical approximations and energy evaluations.

Another quantitative accomplishment is explaining the 10^8^ seeming discrepancy between the Demon’s optimal computation and a neuron’s optimal computation. This explanation hinges on (i) the assumption that there is a bits/sec requirement arising from communication constraints, (ii) a previously derived bits/joule/spike optimization that is based on the unclocked, asynchronous, approximately Poissonian arrival of pulses onto a neuron, and (iii) on the energy-audit provided here. The bits/sec requirement is juxtaposed with the zero bits/sec limit result of physics and further compounded by the slow information growth, 2^−1^ ln(*N*), compared to cost increases that are directly proportional to *N* itself. That this ratio is so unfavorable is not new, at least for sensing (14), but the huge, quantified and explicated discrepancy for computation seems novel.

Additional novel results of our approach include:

1. A precise definition of computation that has meaning both inside and outside of neuroscience, including an explicit role for energy and an explicit Bayesian inference, estimation;
2. An energy audit of the human brain with the relevant partitioning of function; The resulting audit reveals
3. Computational costs are less than 0.2 watts total; in other words, for the average neuron, computation consumes 1.1 · 10^−11^ W whereas GM communication consumes 1.8 · 10^−10^ W (a 16-fold change).
4. Contrary to a reoccurring assumption in discussions of sparse coding, doubling the average firing rate does not double the total signaling costs. Resting potential costs are unaffected by such a doubling, and they account for nearly two-thirds of the gray matter costs at 1.6 Hz. The primary motivations for this energy-audit are calculations of the optimal bits/J which implies the bits/sec. The general, earlier optimization result of (31) is specialized to the current analysis to produce the optimal *N*, the average number of synaptic activations needed to reach threshold.
5. Using the bits/J formulation, the value of *N* ≈2500 inferred in the audit is also the value needed to optimize the bits/J.
6. Using these results and an additional corollary produces the MSE of a neuron’s estimate of its latent variable, and this error decreases in proportion to the energy devoted to computation, 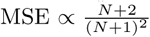.
7. Using the Lindley-Shannon-Bayes valued computations, the values derived in the audit and using the optimal *N*, the computational efficiency is 6.5 · 10^11^ bits/joule per neuron. This optimization also implies the unknown but implicitly constraining communication bit/rate, ca. 4.6 bits/IPI/neuron (7.4 bits/sec/neuron at 1.6 Hz).
8. As part of the information calculation, noise sources are quantitatively compared. The dominating noise-source is inherent in the signal itself; thermal noise is easily ignored while shot-noise is at the threshold level of ignorable.

A common question is, how do we know that the information measure and the definition of computation being used are the right ones? The answer is in two parts: (i) there is not now, nor will there ever be, a provably correct measure or definition. However, (ii) if a chosen measure and definition lead to the optimization of a sensible function in the context of Darwinian evolution, then these definitions are (a) useful, and (b), this utility is justification enough. Thus the claim here is that the measure and the definition being employed show their utility in the Darwinian context of optimized energy use.

### The human brain energy audit compared to the rodent

The per neuron values here are relatively close to those obtained by Herculano-Houzel (39). Her value for the gray matter energy-use of human cortex is 1.32 · 10^−8^ *µ*mol of glucose per neuron per minute, which converts to 2.26 · 10^−10^ W/neuron in terms of ATP. Our value is 1.94 · 10^−10^ W/neuron (Table S3). This small, 16% difference is not surprising since she uses the older glucose values of slightly more than 20 W per brain, and we use her regional brain weight values and cell counts.

The top-down part of the audit can do no more than limit the total ATP available among the defined uses of ATP. Except for this contribution, the top-down calculations are of no use in calculating computational energy use. This is due to the variance of such a top-down calculation since, for the average cortical neuron of the average human, the variance will always exceed the average energy expended for computation. Thus one must rely on bottom-up calculations, and here we look to the landmark work of Attwell and Laughlin (35).

Staying as close to (35) as sensible, newer research is used (e.g., for conversion of glucose to ATP (46) and for the overlapping Na-K conductances of the AP (45)). Species differences also create unavoidable discrepancies, including average firing rate, the fraction of the time that the glutamate-primed NMDARs are voltage-activated, and, more importantly, the surface area of rat axons vs human axons.

Most fundamental for us is the difference in partitioning. Our partitioning begins with the definition of computation and then is further refined by the tripartite distinctions of energy costs: time-proportional, AP-dependent, and failure-rate modified. We acknowledge that this partitioning of energy consumption is at variance with that used in Levy and Baxter’s axon calculations in addition to Attwell and Laughlin’s work. Estimating the cost of *Other* is problematic. The distinctions here require a subpartitioning of *Other* between communication, computation, and the pair synaptic modification and maintanence. But these categories too must be tripartite partitioned. Because computation takes so little of the total energy, only a negligible fraction of *Other* adds to the computational term. More importantly, there is the cost of synaptic modification, including metabotropic receptor activation and postsynaptically activated kinases, which do not fall within the present definition of computation but are activity dependent costs.

### General relevance of Results

#### Outside of neuroscience

Because there is some interest e.g., (47, 48) outside of neuroscience to reproduce neurally mediated cognition on a limited energy budget, the energy-audit here brings an increased specificity to a comparison between the evolved biological vs the human engineered. In particular, engineers often tout brain function as consuming energy at what they consider a modest 20 W given the difficulty they have in reproducing human cognition. Here we provide a more precise set of comparisons. Our computation can be compared to the job performed by the central processing unit. Communication has it’s two major forms defined here, axonal costs and presynaptic functions, which must be compared to communication into and out of memories plus the communication of clock pulses. Perhaps maintenance can be compared to memory refresh costs. However, comparing power conversion loss by a computer to the heat generation of intermediary metabolism is challengeable since heating is fundamental to mammalian performance. A better comparison might be between the cost of cooling a computer and the biological heating cost.

#### Inside neuroscience

Although the primary goal of the energy audit is an estimate of the cost of computation *per se*, the audit also illuminates the relative energetic costs of various neural functions. Notably for humans, the audit reveals that axonal resting potential costs, also called leak, are greater than the firing-rate costs, which seems somewhat surprising. This axonal resting expense is directly proportional to the leak conductance and axonal surface area. Thus, of all the parameters, these two might benefit the most from better empirical data. Regarding these large, leak-associated costs, two additional points seem relevant.

First, regarding fMRI studies that measure regional brain use, the small increases of oxygen consumption over baseline consumption (49) is consistent with the high, continuous cost of axonal leak.

Second, arguing from her data and data of other studies (39), Herculano-Houzel presents the intriguing hypothesis that average glucose consumption per cortical neuron per minute is constant across mammalian species. Qualitatively, this idea is consistent with the increase in neuron numbers along with the decrease of firing rates found in humans vs rats. However, it seems that the hypothesis can only be quantitatively correct if axonal leak-conductance in humans is much lower than in animals with smaller brains and presumably shorter axons of smaller diameters. This topic deserves more detailed exploration.

Hopefully the work here motivates further empirical work, especially using primates, to improve the energy-audit and the calculations that ensue. Such empirical work includes better surface area measurements and a better idea about the NMDAR off-rate time constant. Finally, going beyond the average neuron, perhaps someday there will be energy-audits matched with the neurophysiology of identified cell types.

## Materials and Methods

**Proofs**. The proof of *lemma 2a* is just a textbook change of variable from one density to another (50) where 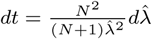 to prove *corollary 1* and the first equality of *lemma 2b*, use *2a* to calculate the appropriate conditional moments, which Mathematica obliges; to prove the second equality of *2b*, use lemma 1 to calculate the indicated conditional moment.

### Parameterizing the marginal prior p(λ)

As derived from first principles in Levy, Berger, and Sungkar (31), the only known, consistent marginal prior of the latent RV is 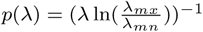 where the bounds of the range of this RV, and thus its normalizing constant, are the subject of empirical observations and the required definition *λ ∈* (0 *< λ*_*mn*_ *< λ* _*mx*_ *< ∞*). Recall that *λ* is the rate of activations of 10^4^ input lines undergoing a 25% success rate when activated. From the energy-audit, use the 1.6 Hz average firing rate. Then *E*[Λ], the mean marginal input firing rate scaled by a 3/4 failure rate is 4000 events/sec (10^4^ · 1.6 · 0.25). Then supposing that the rate of spontaneous release is 1 Hz over these 10^4^ synapes, *λ*_*mn*_ = 1. With one unknown in one equation, 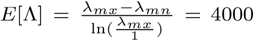, Mathematica produces *λ*_*mx*_ ≈ 42643, and the prior is fully parameterized.

### Adjusting the bit-rate calculation for multiple IPIs per decision-making interval (DMI)

The nearly 7 bits per IPI only applies to a neuron’s first IPI. Later spikes are worth considerably less using the current simplistic model of a fixed threshold. Using the prior distribution, a neuron does not fire 37% of the time within the 625 msec DMI while 63% of the time, a neuron fires one or more times in the DMI. As a crude approximation, suppose 26% of the time a neuron fires two or more times, 8% of the time a third spike is produced in an DMI, and 3% of the time a fourth spike is produced. Thus the average number of spikes per DMI is appropriately one. Using a simplistic model with a fixed threshold of N, the bit values of the later spikes are quite small. The value of the second through fourth spikes are 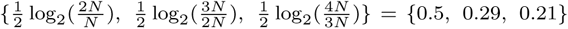 respectively. The weighted value ac^*c*^ounting for all spikes is then ca. 4.6 bits/DMI rather than the almost 7 bits of the first IPI.

### Shot-noise has a nearly negligible effect on bit-rate

As measured in the biophysical simulations (23), the most deleterious degradation of a neuron’s computation arises, not from thermal noise or shot-noise (24), but from the neuron’s input signal itself. Here is a calculation consistent with this biophysical observation.

Using stochastic NaV 1.2 and NaV 1.6 channels in a biophysical model of a rat pyramidal neuron, it is possible to observe shot-noise and to estimate the number of such channels that are activated at threshold. With relatively slow depolarization, there are less than 250 channels on when threshold is reached, and this number of channels seems to contribute less than 1.6 mV (see Fig 5 in (23)). Thus modeling channel activation as a Poisson process with rate 250 and individual amplitudes of 6.4 *µ*V, Campbell’s theorem (51) produces the variance; this variance is less than 250 · (6.4 ·10^−6^)^2^ = 1.6 ·10^−9^. The same calculation for the input excitation yields a variance of 2500 · (6.4 ·10^−6^)^2^ = 1.6 · 10^−8^. Then, the net variance is represented by multiplying the drift variance by something that increases it less than 10%, say 1.09. This nine percent greater variance reduces the information gain by log(1.09) ≈ 0.12 bits.

### Numerically-based optimization calculations

Optimizing the bits/joule equation uses Mathematica. As threshold and the average synaptic event are continuous variables, the calculated *N*, the average number of events per IPI, is also a continuous variable. Then to optimize, we take the derivative, *dN*, of the single neuron, single IPI bit/J formulation. Then setting the numerator of this derivative equal to zero, we solve for *N* using Mathematica’s NSolve. We also examine the optimal *N* for different values of energy-use. Although the N-optimization requires solving a transcendental equation, the optimal *N* is nearly a linear function of the ratio of AP-communication costs vs computational costs. Specifically, 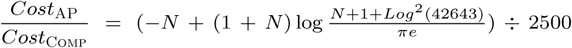, an expression that makes as plain as possible the required relationship between the relevant energy consumption terms and the assumption that *N* = 2500.

### Partitioning glucose by region and by metabolic fate

This section explains the top-down calculations of Table 1. The glucose-uptake values combine the regional uptakes, reported in terms of per 100 gm of tissue from Graham et al. (38) as copied into our Table S1 along with the reported regional masses from Azevedo et al. (52). We choose this uptake study because of its use of the [^11^C]glucose tracer and its straightforward application to obtain regional net glucose uptakes. Multiplying regional masses by uptake values, and converting to appropriate units as in Table S1, yields the first “Watts” column of Table 1. These glucose-watts are calculated using 2.8 MJ/mol (53). The regional uptakes are combined to produce the brain total as illustrated in Fig S1.

Following the flow diagram of Fig S1, next we remove the nonoxidized glucose from regional and total uptakes. We use an oxygen-glucose index (OGI) value of 5.3 (out of 6 possible oxygen molecules per one glucose molecule). We assume the OGI is constant across regions and that we can ignore other, non-CO_2_ carbons that enter and leave the brain. Thus, these simple glucose-watts are split into oxidized and non-oxidized as produced in Table 1 and illustrated in Fig S1.

As the energy source, the oxidized glucose is then partitioned into two different metabolic fates: heating and ATP. Again we assume this process is constant across regions and that the brain does not differ too much from other regions which have been studied in greater depth. The biological conversion is calculated using Nath’s torsional mechanism, which yields 37 ATP molecules per molecule of glucose and 36,000 J/mol of ATP at 37^*°*^ C.

The definition of computation and its distinction from communication produces the partitioning here that differs from earlier work. As opposed to the earlier neuroscientifically oriented thinking (35), the motivating perspective here is computational function. That is, computation and its cost are specifically identified with the charging and discharging of the dendrosomatic plasma membrane. For this reason, we emphasize the reset and charging as opposed to resting potential for the dendrosomatic membrane. A second distinction is “synaptic” energy-use. The purely neuroscientific perspective follows traditional morphogical distinctions by considering synaptic costs as a whole, pre- plus postsynaptic parts. This perspective differs from the distinction made here where pre- and postsynaptic costs are separated. Thus, once computation is defined, it is clear that presynaptic function is, inclusively, just the endpoint of long-distance communication. To put it another way, each portion of a neuron needs to be accounted for, but each portion can only be counted once.

More details concerning the partitioning here versus earlier work are found in the Supplement.

### Computation Costs

Our “on average” neuron begins at its reset voltage and then is driven to a threshold of −50 mV and then once again resets to its nominal resting potential of −66 mV. Between reset and threshold, the neuron is presumed to be under constant synaptic bombardment with its membrane potential, *V*_*m*_, constantly changing. To simplify calculations, we work with an approximated average *V*_*m*_, *V*_*ave*_ of −55 mV; this approximation assumes *V*_*m*_ spends more time near threshold than reset. (Arguably the membrane potential near a synapse which is distant from the soma is a couple of mVs more depolarized than the somatic membrane voltage, but this is ignored.) To determine the cost of AMPAR computation, we use the ion preference ratios calculated from the reversal potential and use the total conductance to obtain a Na^+^ conductance of114.5 pS per 200pS AMPAR synapse as seen in Table S4. (The ion-preference ratios used for the calculations in Table S4 are calculated from the reported reversal potential value of −7 mV (54) and the individual driving forces at this potential, −90 − (−7) = −83 *mV* for K^+^ and 55 − (−7) = 62 *mV* for Na^+^.) Multiplying the conductance by the difference between the Na^+^ Nernst potential and the average membrane potential (*V*_*Na,Nern*_ − *V*_*ave*_) yields a current of 12.5 pA per synapse. Multiplying this current by the SA duration, converts the current to coulombs per synaptic activation, and dividing this by Faraday’s constant gives us the moles of Na^+^ that have entered per synaptic activation. Since 1 ATP molecule is required to pump out 3 Na^+^ molecules, dividing by 3 and multiplying by the average firing rate yields 8.35 ·10^−20^ mols-ATP/synapse/sec. Multiplying by the total number of synapses adjusted by the success rate (0.25 1.5 ·10^14^ synapses) implies the rate of energy consumption is 0.113 W for AMPAR computation. When NMDARs are taken into account, the total computational cost is 0.17 W (assuming that NMDARs average conductance is half as much as AMPAR’s).

Table S4 lists the excitatory ion-fluxes mediated by AMPARs and NMDARs. The cost of the AMPAR ion fluxes is straightforward. The cost of NMDARs ion fluxes depends on the off-rate time constant as well as the average firing rate. That is, if this off-rate time constant is as fast as 100 msec and the IPI between firings of the postsynaptic neuron is 400 msec or more (such as the 625 msec interval that comes from the 1.6 Hz frequency used in the following calculations), then most glutamate-primed NMDARs will not be voltage activated. Thus, in contrast to the rat where the AMPAR and NMDAR fluxes are assumed to be equal, here we assume the ion-fluxes mediated by NMDARs are half that of the AMPARs and multiply the AMPAR cost by 1.5 to obtain the final values in Table S4.

The spike-generator contributes both to computation and to communication; fortunately, its energetic cost is so small that it can be ignored.

### Communication Costs

Table S5 provides an overview of the communication calculations, which are broken down into Resting Potential Costs, Action Potential Costs, and Presynaptic Costs. The following sections explain these calculations, working towards greater and greater detail.

In general, the results for communication costs are built on less than ideal measurements requiring large extrapolations; here are some examples. There does not seem to be any usable primate, much less human data. The proper way to determine surface area is with line-intersection counts, not point counts, and such counts require identification of almost all structures. As the reader will note in the supplement, use of mouse axon diameters produces much larger surface areas, thus raising communication costs and decreasing the energy available for computation and *Other*. Likewise, copying recent values used in the biophysical literature for axon resting resistance (a rather difficult parameter to measure, especially for the small axons of interest here) also greatly increases the cost of communication compared to the values that we used (45, 55).

### Resting Potential Costs

The cost of the resting potential itself is simply viewed as the result of unequal but opposing Na^+^ and K^+^ conductances. If other ions contribute, we just assume that their energetic costs eventually translate into Na^+^ and K^+^ gradients. The axonal resting conductance uses a value from biophysical simulations of rat pyramidal neurons, although higher values are not uncommon (e.g., (45, 56)). Resting potential costs of axons (including axonal boutons) assume a resting, passive resistance of 30 kΩ cm^2^ and a membrane surface area of 21.8 · 10^6^ cm^2^ (see Table S6), producing a total conductance of 727 S. The driving voltage for each ion is determined by subtracting the appropriate Nernst potential from the assumed resting membrane potential of −66 mV. Using Nernst potentials of +55 mV and −90 mV for Na^+^ and K^+^ resp., we just assume currents are equal and opposite at equilibrium. Thus, conductance ratios are calculated from the equilibrium condition: −24 mV *·g*_*K*_ = −121 mV *·g*_*Na*_; implying *g*_*K*_ = 5.04 *g*_*Na*_; and further implying 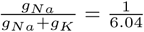. The Na^+^-conductance times the driving voltage yields the 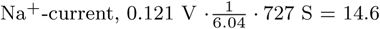 A. Divide this result by Faraday’s constant to find the total *Na*^+^ influx, and then divide by 3 to obtain the number of ATPs required to pump out this influx, 5.03 · 10^−5^ molATP/s. Multiplying this number by 36,000 J/molATP yields 1.81 W, the resting potential cost.

Plasma membrane leak is a major energy expenditure in both the calculations here (66% of gray matter communication costs) and in the Attwell and Laughlin calculations (13% of signaling-related ATP consumption). The differences in these percentages arise from rather different interpretations of a functioning neuron and of the meaning of certain measurements. Here there is an important distinction between the cost of reset vs the cost of resting potentials: the resting potential cost is entirely axonal and essentially continuous across time. On the other hand, the cost of resetting synaptic depolarization applies only to the dendrosomatic portion of a neuron, and this portion of a neuron is under constant synaptic bombardment. Thus resting potential in this portion of a neuron is quite transient. In contrast to the calculations here, the rat calculation uses the somatic measurement, which we contend is primarily evaluating dendritic conductance to ground.

### Action Potential Costs

Action potential costs are calculated from Na^+^ pumping costs as delineated in Table S5. The coulombs to charge a 110 mV action potential over the entire non-bouton axon starts with the product of the total GM axonal capacitance, 14.6 F, the peak voltage, and the firing rate, 1.6 Hz; i.e., 14.6 ·0.11 ·1.6 = 2.57 amps. To account for the neutralized currents observed by Hallerman et al. (45), multiply the previous result by 2.28, yielding 5.86 A.

Bouton costs, although clearly part of an axon, are calculated separate from the axon. As will be detailed later, our approximation of surface areas treats all presynaptic structures as *bouton terminaux*, and rather than assume tapering for impedance matching purposes, presume an abrupt transition of diameters. Importantly, we assume that a bouton mediates a calcium spike and that this spike only requires a 0.02 V depolarization to be activated. Altogether, the rate of *Na*^+^ coulomb charging for boutons is 6.34 F ·0.02 V ·1.6 Hz = 0.20 A.

The sum of axonal spike costs and bouton chargings is used to determine the Na^+^ that needs pumping. Thus, dividing the total current by Faraday’s constant converts coulombs per sec to mols of charge per sec, and this calculation yields a Na^+^ flux of 6.3 · 10^−5^ molNa^+^per sec. Dividing by three converts to ATP mol/sec, and multiplying this value by Nath’s 36,000 J/molATP yields the total action potential cost of 0.75 W.

As noted earlier, the WMAP costs are required. To approximate this value, assume that the ratio of GMAP cost to total GM axonal cost equals the ratio of WMAP cost to the total WM cost. Thus, 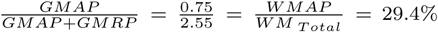 then with *WM*_*T otal*_ = 1.85 W, WMAP = 0.54 W.

Since some portion of *Other* is likely AP-dependent, we scale the 0.17 W cost of *Other* in the same proportion as the GM communication costs scale for APs vs APs plus rest potential (where APs include presynaptic AP costs): 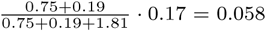 W.

### Presynaptic AP Costs

The presynaptic transmitter-associated costs are mostly based on the values of Attwell and Laughlin (35) and of Howarth et al. (43). The assumptions include an assumed 25% success rate of vesicular release for each cortical spike (2.4 · 10^14^ spikes/sec under the 1.6 Hz and 1.5 · 10^14^ synapses assumptions). However, in contrast to Howarth et al. (43), which uses a number supported by observations in calyx of Held (57) and in cell cultures (58), the observations of Stevens and Wang (44) in CA1 hippocampal pyramidal neurons indicate that the same calcium influx occurs for both synaptic successes and failures. Because adult hippocampal synapses seem a better model of cerebral cortical synapses then calyx or tissue culture synapses, we use the hippocampal observations. Therefore, the 1.6 Hz firing rate produces a Ca^2+^ cost that is more than 8-fold greater than the cost of vesicle release events (VR events, Table S5). The Ca^2+^ influx per action potential is 1.2 · 10^4^ Ca^2+^/vesicle, and assuming 1 ATP is required to pump out each Ca^2+^, the Ca^2+^ cost is 1.2 · 0^4^ ATPs/vesicle. Multiplying this by 2.4 · 10^14^ APs/sec for the gray matter yields a total presynaptic Ca^2+^ cost of 0.17 W.

The cost per vesicle release is determined by adding the packaging and processing costs and then multiplying by the number of glutamate molecules per vesicle as in (35) and (43). Adding the cost of membrane fusion and endocytosis yields a total of 5,740 ATPs/vesicle (43). This value is multiplied by the VR events per second and divided by Avogadro’s number to obtain 5.7 · 10^−7^ ATPmol/sec. Converting to watts yields a presynaptic transmitter release cost of 0.02 W and a total presynaptic cost of 0.19 W for the GM.

### Axonal and presynaptic surface area

Surface areas of axons and their associated presynaptic structures are critical to the estimation of gray matter communication costs. Alas, the lack of human data forces several bold extrapolations. Fortunately, some EM volume-fraction observations in other species and one well-quantified light microscopic (LM) study in cats help to constrain or serve as a check on our assumptions.

### Synapse counts

Both computation and communication costs depend on the number of cortical synapses. For the approach taken here, computational costs scale in a one-to-one ratio to synaptic counts while communication costs scale proportionally, but with a smaller proportionality constant.

The calculations use the Danish group’s synapse counts of 1.5 · 10^14^ (59). The alternative to the numbers used here report an 80% larger value (60); however, their human tissue comes from nominally non-epileptic tissue from severely epileptic patients. Since the incredibly epileptic tissue is likely to stimulate the nearby nonepileptic tissue at abnormally high firing rates, we find the data’s import questionable.

### Estimation of Surface Areas from Mouse and Rabbit Data

Here volume-fraction data are used to estimate axon and presynaptic surface areas. As far as we know, there are two journal-published, quantitative EM studies of cerebral cortex that are suitable for our purposes: one in rabbit (61) and one in mouse (62). (Although structural identifications do not neatly conform to our simplifying cylindrical assumptions, we can still use their data to direct and to check our estimates.)

Chklovski et al. (62) report a 36% volume-fraction for small axons, 15% for boutons, 11% for glia, 12% for other, and 27% for dendrites and spines as read from their graph in their Figure 3. They purposefully conducted their evaluations in tissue that lacked cell bodies and capillaries. Because cortical tissue does contain cell bodies and capillaries, this will produce a small error for the average cortical tissue. More worrisome is the size of “other,” half of which could be very small axons.

The quantification by Schmolke and Schleicher (61) examines the rabbit visual cortex. Their evaluation partitions cortex into two types of tissue: that with vertical dendritic bundling and that which lacks dendritic bundling (they do not seem to report the relative fraction of the two types of cortex, but we assume the tissue without bundling dominates over most of cortex). For boutons and axons respectively, they report volume fraction values within bundles of 17% and 20% and values between bundles of 26% and 29%.

The 30% axonal volume fraction used in Table S6 is a compromise between the (62) value of 36% and the two values from (61). The average of the within bundle and between bundle volume-fractions from (61) is used for boutons. Specifically, the approximated human volume fractions are (i) 22% boutons, (ii) 30% small axons, (iii) 11% glia, (iv) 5% neuronal somata, (v) 3% vasculature, (vi) 29% dendrites, spineheads, and spine-stems, totaling 100%. (It is assumed that standard fixation removes almost all of the physiological extracellular space and, naively, shrinkage/swelling has little relative effect on these values.) The calculations are essentially unaffected by the two conflicting bouton volume fractions since the difference between the two possible calculations is negligible.

Table S6 lists the critical values, the intermediate values for the cylindrical model to fit the data, and finally the implications for the relevant membrane capacitance.

### Cylindrical model approximations for axons and boutons

#### Axons

By making a cylindrical assumption and assuming the average small axon’s diameter is 0.50 *µ*m (radius = 0.25 · 10^−4^ cm), we can estimate the total surface area of these unmyelinated axons using the 30% volume-fraction to calculate the length of an average axon, *L*_*ax*_. The total volume (cm ^3^) occupied by all such axons is *L_ax_* · 1.5 · 10^10^ · *π*(0.25 · 10^−4^)^2^. Dividing this volume by the volume of the GM (632 cm^3^) must equal the volume fraction, 0.3. Solving yields *L*_*ax*_ = 6.44 cm. Then net surface area is calculated using this length, the same diameter and number of neurons, 6.44 · 1.5 · 10^10^ · *π* · 0.5 · 10^−4^ = 1.52 · 10^7^ cm^2^. For an independent calculation of axon length based on LM data, see Supplement.

#### Boutons

The surface area estimates also treat boutons (Btn) as uniform cylinders of a different diameter. Assume that cortical presynaptic structures in humans are no bigger than in any other mammalian species. To determine bouton surface area, assume a bouton diameter (*d*_*pb*_) 1.1 *µm* and height (*h*_*pb*_) 1.0 *µm*. Denote the total number of synapses in the gray matter as *n*_*gm*_ (1.5 · 10^14^). (Note that the cylinder area of interest has only one base.) Then, with the formulation 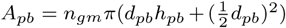, the bouton surface area works out to *A*_*pb*_ = 1.5 · 10^14^*π*(1.1 *µm* · 1.0 *µm* +(0.55 *µm*)^2^) =*·*6.61 · 10^6^ cm^2^. See Tables S6 and S7.

We assume a bouton only accounts for one synapse. However, larger boutons can contact multiple, distinct postsynaptic neurons. Thus the small cylinders, as individual synapses, are an attempt to approximate such presynaptic configurations. See Table S8 for more details and for the effect of overestimating areas.

### Oxidized vs. non-oxidized glucose

Arteriovenous blood differences indicate that insufficient oxygen is consumed to oxidize all the glucose that is taken up by the brain. Supposing glucose is the only energy-source, it takes six O_2_’s for complete oxidation. The calculations use an OGI value of 5.3 (63). Other values from arteriovenous differences are found in the literature (64–66). Even before these blood differences where observed, Raichle’s lab proposed as much as 20% of the glucose is not oxidized (40).

### Glucose to ATP based on Nath’s theory

Table S2 offers the reader a choice between Nath’s torsional conversion mechanism of glucose to ATP (46, 67, 68) versus the conventional conversion to ATP based on Mitchell’s chemiosmotic theory (69). According to Nath, the minimum number of ATP molecules produced per molecule of glucose oxidized is 32, and this includes mitochondrial leak and slip (46). Nath’s calculations are based on free-energy values under physiological conditions. However, his calculations are recent while the standard model has been taught for decades, although not without controversy (70). The standard textbook number for this conversion is 33 ATPs per molecule of glucose before accounting for mitochondrial proton leak and slip. Since leak is often assumed to consume 20% of the energy that might have gone to ATP production in oxidative phosphorylation (35, 71), the Mitchell conversion number is reduced from 33 to 27 molecules of ATP (2 ATPs are produced by glycolysis and 2 by the Krebs cycle, so this 20% reduction only applies to the ATP produced in the electron transport chain).

The other choice given to the reader in Table S2 is the choice between two different firing rates. When the higher firing-rate or the Mitchell mechanism is used, there is no energy available for *Other*. Thus in these cases, the accounting cannot be balanced. In this regard, an energy-allocation for maintenance and synaptic modification (*Other* in Table 1 and 2) is a bare minimum and is just estimated via the guess that its value is equal to the computational cost.

### Other

Here *Other* is not directly calculated. Rather it is matched to computational energy consumption. As noted in Discussion, this category must itself be partitioned into three types of energy consumption. We assume that *Other* partitions in direct proportion to energy-use elsewhere. Fortunately, our information calculations will hold if energy is exchanged between categories with similar dependencies. For example, *Other* needs to be partitioned between the two types of communication costs (APs vs resting potential), costs arising from postsynaptic depolarization, and the costs arising from metabotropic activations and synaptic plasticity. See Supplement for further explications of *Other* regarding partitioning and firing rate dependency.

## ACKNOWLEDGMENTS

The authors are grateful for comments and suggestions of earlier versions provided by Costa Colbert, Robert Baxter, Sunil Nath, and David Attwell.

