## Supplemental Information for "Computation in the human cerebral cortex uses less than 0.2 watts yet this great expense is optimal when considering communication costs"

PNAS
www.pnas.org

### Supplementary Information for

#### Your main manuscript title

**William Levy, Victoria Calvert**

**William Levy**
****

##### This PDF file includes:

Supplementary text
Fig. S1
Tables S1 to S8
SI References

#### Supporting Information Text

**Selecting, adjusting, and comments on literature values for glucose uptake .** In the literature, there are various uncertainties and incompatibilities which require compromises and approximations. The subcortical distinctions made by Azevedo et al. (1) and Graham et al. (2) are different. That is, the purely anatomical study lumps together the striatum, thalamus, colliculus, hypothalamus, pons and medulla while the reported [$^{11}$C]glucose measurements only include the individual subcortical regions caudate, putamen and thalamus. Assuming these subcortical forebrain regions account for much of the weight of what Table 1 and Table S1 labels as "other regions", and assuming that the unmeasured brain regions are not too different in glucose uptake, we just use a single value and multiply by the weights of the subcortical regions given by Azevedo et al. (1).

The [$^{11}$C]glucose uptake by the choroid plexus and brain capillaries is assumed to be negligible. The ventricular weight is inferred from Azevedo et al. (1) to produce their total of 1510 g.

Another approximation is required due to the non-uniformity of brain size. The representative brain mass values available are for an average male brain. Incompatibly, the regional glucose uptake values from Graham et al. (2) are averages from six females and four males with no statements about sex differences. Due to this sex heterogeneity there will be large total brain weight heterogeneity and a lower average brain mass per subject than the 1510 g value. Thus some scaling or conversion is needed. To remain consistent, the Graham et al. regional uptake values are scaled by the Azevedo et al. regional brain weights. Table S1 details these calculations. As a result, instead of a total brain uptake rate of 6.48 $\mu$ mol/sec the summed regional uptake rates yields 6.05 $\mu$ mol/sec.

The OGI number is problematic in terms of accuracy and reproducibility. One group with multiple publications on the topic reports an OGI value of slightly more than 5.2 (3). Other A-V studies favor a higher OGI (ca. 5.6, (4)), but several of these studies also favor decreasing the value of glucose uptake by virtue of the net efflux of non-$CO_2$ carbons. Supposing the larger OGI is correct and supposing that we are allowed to ignore non-$CO_2$ carbon efflux, such an increase in oxidized carbons could yield an additional ca. 0.14 ATP-watt available to gray matter.

As Fig. 2 in the main text and Fig. S1 show, we do not bother to subdivide the energy-use of the white matter beyond its total ATP-watts. In contrast, gray matter is further subdivided. Specifically, separate bottom up calculations are used to determine communication and computational energy consumption (see below).

**Conversion of glucose to ATP.** There are three problems with the textbook calculations and previous brain calculations of glucose to ATP conversion: such calculations are based on room temperature free-energies, the mitochondrial leak value is based on a maximal leak value from non-neuronal tissue, and the chemiosmotic hypothesis ignores slip in redox pumps. Slip would further reduce the chemiosmotic ATP values by 10% (Nath, personal communication). However, Nath's novel mechanism does not require any ATP downgrades. (He identifies the neutral form of succinic acid as mediating leak. This form can penetrate the cristal membrane as the dianionic form, creating slip while the succinate monoanion is the motive form. Thus, both leak and slip are accounted for (5).) Here, however, we do not attempt to account for the issue of slip in our chemiosmotic calculations. Thus, the choice offered for ATP production in the gray matter in Table S2 is 3.09 W (Nath) or 2.61 W (Mitchell). Both of these calculations use the conversion factor of 36 kJ/molATP (5, 6) rather than the room-temperature value typically used.

Although we consider Nath's torsional mechanism to be a more accurate depiction of ATP production than the chemiosmotic mechanism, we recognize that this newer mechanism is not directly informed by brain mitochondrial studies. That is, brain tissue has different uncoupling proteins which could potentially alter the amount of ATP produced per mole of glucose. Of course, different species have different issues concerning thermoregulation and thus may differ in amount of leak (see for example (7)). Such studies argue that larger animals are more efficient in regard to mitochondrial production of ATP. Thus the conversion values of glucose to ATP used for rats are plausibly lower than for humans.

##### Other contrasting estimates with earlier results in the literature.

***Postsynaptic ionotropic costs are different.*** Our postsynaptic costs for AMPAR activation are about half of Attwell and Laughlin's ($77 \cdot 10^6$ ATP/action potential/neuron vs $134 \cdot 10^6$ ATP/action potential/neuron) (8). The difference mostly arises from their use of an outlier value for synaptic conductance that is doubted by the authors themselves as cited in Attwell and Laughlin. Biophysical simulations encourage us to discard the outlier value.

The biophysical simulations in Singh and Levy (9), which use the consensus layer 5 prefrontal pyramidal neuron found in a variety of biophysical models, support the lower postsynaptic activation costs used here. Such simulations use 200 pS AMPAR-conductances per synapse and no NMDARs. Under these conditions, it typically takes 250 to 700 synaptic activations to fire the neuron. Such simulations did not include inhibition. Upgrading the model to include inhibition, as is done here and as occurs in the several articles that consider balanced inhibition (e.g., (10, 11)), is required to be consistent with the estimate of 2500 synaptic activations per output spike. That is, inhibition is implicitly incorporated into our cost calculations by virtue of increasing the number of synaptic activations needed to reach threshold.

The same amount of excitatory synaptic activations (ca. 2500 input activations per pulse out) can be true for both humans and rats. That is, although the dendritic surface area of a human cortical neuron is greater than a rat's, a lower rate of inhibition in the human can compensate so that the same amount of excitatory synaptic activation is required to fire either neuron.

***Sensitivity to axon and presynaptic assumptions.*** Gray matter communication costs are rather large and are directly a function of axonal and presynaptic surface areas. Therefore, it is worth revealing the sensitivity of the calculations to the assumptions going into these surface area calculations.

*Axons*: Fixing the volume-fraction at 30%, and varying axon diameter changes the values of surface area and axon length. Table S7 shows the relationship between these parameters as well as their affect on aspects of communication costs. Mouse data motivates much narrower axons but not smaller than 0.25 $\mu$m (12). As the smallest possible diameter, we choose the 0.28 $\mu m$ mouse-inspired diameter (13).

Merely changing diameter from 0.5 to 0.4 $\mu$m increases the axonal contribution to total communication costs by 0.5 W. Clearly the mouse diameter increases energy-use well beyond that which is available based on top-down calculations.

*Boutons*: As noted before, the cylindrical assumption for boutons is crude. Table S8 illustrates the sensitivity of our bouton size assumptions and, by extension, our bouton capacitance values relative to volume fractions. In these calculations, the different volume fractions are a necessary result of varying the bouton dimensions.

***Reconciling the cost of Other.*** First, it must be said that the catchall partition labeled *Other* has never been properly measured for an adult animal as far as we know. More to the point here however, is that our perspective on what *Other* should include differs somewhat from earlier work.

What follows explains our rejection of previous estimations of the ATP-use attributed to the catchall and unpartitoned "*Other*" category of energy consumption. In the rodent audit, the energy consumed by *Other* is based on published research that: (i) mathematically differences ATP-use before and after inhibition of the Na-K ATPase pump and that (ii) implicitly but necessarily assumes that removing the $Na^+$ gradient will not increase other forms of ATP-use. However, the literature argues otherwise. In general, cardiac glycosides such as ouabain, will accelerate several forms of ATP consumption.

Ouabain causes depolarization because of leak conductance, and it also causes transmitter release (14–17) and in particular quantal transmitter release (18). Quantal transmitter release implies (i) the ATP-consuming processes of vesicle recycling and re-loading, (ii) postsynaptic activation of GTP-consuming metabotropic receptors, and (iii) postsynaptic activation of calcium-conducting NMDARs, which will activate various postsynaptic, ATP-consuming kinases. Moreover, in addition to the vesicle recycling, metabotropic, and kinase costs, there are both pre- and postsynaptic calcium pumping costs. That is, poisoning the Na-K pump leads to increase of $[Ca^{2+}]_{in}$ as does the NMDAR activation. Raising levels of internal calcium in turn activates one or more of at least three types of $Ca^{2+}$-ATPases: those in the plasma membrane, those associated with sarco- and endoplasmic reticula, and a mitochondrial accumulator (19–21). Any other kind of poisoning experiment that causes depolarization will produce similar increased demands for ATP. Finally, there are those (not Attwell et al.) that assume blocking action potentials will remove communication costs, but our leak calculations refute this idea.

***Comparison to a class of completely quantified axons.*** Concerning axon lengths, there is one LM study that provides data indicating the reasonable nature of the length estimates obtained above. In particular, there are axonal length data for cat L2/3 pyramidal cells using 30 completely stained axons. These data imply an axonal length of 4 cm per neuron (22), but as LM measurements, they will incorporate terminal boutons. For comparison to the estimates here, we combine axonal boutons and the axonal lengths without terminal boutons. Assume there are ten thousand boutons per neuron, each $1.1 \cdot 10^{-4}$ cm in length. Using a volume fraction for small axons of exactly 30%, our unadorned axon length is 6.44 cm per neuron. Adding the bouton lengths yields 7.44 cm per neuron. Thus we are predicting that LM quantification of the average human pyramidal neuron's axon is about 86% longer than the cat axon. More details on the derivations and parametric sensitivities are found in Supplement including Tables S6-S8.

#### Probability and entropy approximations.

***Initial development of approximations.*** There are two approximations needed to calculate a valued Lindley-Shannon information rate, $h(\hat{\Lambda}) - h(\hat{\Lambda}|\Lambda)$. After an exact integration by Mathematica, the first approximation follows. Specifically,

$p(\hat{\lambda}) = \int_{\lambda_{mn}}^{\lambda_{mx}} p(\lambda)p(\hat{\lambda}|\lambda)d\lambda =$

$(\hat{\lambda}\ln(\frac{\lambda_{mx}}{\lambda_{mn}}))^{-1} \cdot \frac{1}{2}(\text{erf}\left(\frac{N\lambda_{mx}-(N+1)\hat{\lambda}}{\sqrt{2(N+1)\hat{\lambda}\lambda_{mx}}}\right) - \text{erf}\left(\frac{N\lambda_{mn}+(N+1)\hat{\lambda}}{\sqrt{2(N+1)\hat{\lambda}\lambda_{mn}}}\right) + e^{2N}\left(\text{erf}\left(\frac{N\lambda_{mn}-(N+1)\hat{\lambda}}{\sqrt{2(N+1)\lambda_{mn}\hat{\lambda}}}\right) - \text{erf}\left(\frac{N\lambda_{mx}+(N+1)\hat{\lambda}}{\sqrt{2(N+1)\lambda_{mx}\hat{\lambda}}}\right)\right))$

$\approx (\hat{\lambda}\ln(\frac{\lambda_{mx}}{\lambda_{mn}}))^{-1} = p(\lambda)$, and therefore,

$h(\hat{\Lambda}) \approx h(\Lambda)$.

Remarks: (i) Mathematica performs the exact integration when one temporarily substitutes $m$ for $(N+1)$ and specifies the appropriate assumptions. (ii) With $N = 2500$, and a naive use of Mathematica, $p(\hat{\lambda})$ approximations of $p(\lambda)$ appears exact when running at the default precision of Mathematica. The erf terms in $p(\hat{\lambda})$ combine to a value of two, which is the exact value needed (since this value is multiplied by one-half).

The second approximation concerns the conditional differential entropies. Recall from Results, $p(\hat{\lambda}|\lambda) = \sqrt{N+1}(2\pi\lambda\hat{\lambda})^{-1/2}\exp(-\frac{\lambda N^2}{2(N+1)\hat{\lambda}} - \frac{\hat{\lambda}(N+1)}{2\lambda} + N)$. Thus in bits and according to Mathematica, $-h(\hat{\Lambda}|\lambda) = (\ln(2)^{-1}(\frac{1}{2}\ln(2\pi e(N+1)^2/N) - \ln(\lambda) + \frac{\exp(2N)\sqrt{N}BesselK^{(1,0)}[-\frac{1}{2},2N]}{\sqrt{\pi}})$. Before valuing this entropy to yield the second approximation, there is a simplification that obviates the need for a third approximation. When the expectation $h(\hat{\Lambda}|\Lambda) = \int p(\lambda)h(\hat{\Lambda}|\Lambda=\lambda)d\lambda$ is taken, the term $E[\ln(\Lambda)]$ appears. However, this same term of opposite appears in the marginal differential entropy $h(\Lambda)$, so they combine to zero. Then with $\frac{\exp(2\cdot 2500)\sqrt{2500}BesselK^{(1,0)}[-\frac{1}{2},2\cdot 2500]}{\sqrt{\pi}} < 10^{-3} \approx 0$, we have the second approximation, and this approximation improves as $N$ increases. The information gain in continuous time (per sec) then is $h(\hat{\Lambda}) - h(\hat{\Lambda}|\Lambda) \approx \log_2(\ln(\frac{\hat{\lambda}_{mx}}{\hat{\lambda}_{mn}})) + \frac{1}{2}\log_2(2\pi e(N+1)^2/N)$. With $\frac{1}{2}\log_2(2\pi e) \approx 2.05$, with $N = 2500$, , $\frac{1}{2}\log_2(2501^2/2500) \approx 5.64$, and $\log_2(\ln(\frac{\hat{\lambda}_{mx}}{\hat{\lambda}_{mn}})) = \log_2(\ln(42643)) \approx 3.41$, the resulting bit rate is 11.1 bits/sec. Given the assumed average firing rate of 1.6 Hz, this evaluates to 6.94 bits/IPI.

***Evaluating the marginal approximation.*** Just below is the Mathematica analysis that investigates the error of the second approximation above. As the Mathematica output demonstrates, the distribution is off by about 1.5 parts in 10,000 accruing from furthest ends of the marginal density as accompanying plots illustrate.

(*The conditional density *)

In[768]:=
```
phatλgivλ =
  Sqrt[(N + 1) / (2 * π * hatλ * λ)] * Exp[N - λ * N^2 / (2 * (N + 1) * hatλ) - hatλ * (N + 1) / (2 * λ)]
```

Out[768]=
$$\frac{e^{N-\frac{\text{hat}\lambda\,(1+N)}{2\,\lambda}-\frac{N^2\,\lambda}{2\,\text{hat}\lambda\,(1+N)}}\sqrt{\frac{1+N}{\text{hat}\lambda\,\lambda}}}{\sqrt{2\,\pi}}$$

In[769]:=
```
Integrate[phatλgivλ, {hatλ, 0, Infinity}, Assumptions → {λ > 0, N > 1}]
```

Out[769]= 1

(* The marginal density of the latent variable *)

In[770]:=
```
pλ = (λ * Log[λmx / λmn])^(-1)
```

Out[770]=
$$\frac{1}{\lambda\,\text{Log}\left[\frac{\lambda\text{mx}}{\lambda\text{mn}}\right]}$$

In[771]:=
```
Integrate[pλ, {λ, λmn, λmx}, Assumptions → 0 < λmn < λmx]
```

Out[771]= 1

(* The joint density *)

In[772]:=
```
PowerExpand[pλ * phatλgivλ]
```

Out[772]=
$$\frac{e^{N-\frac{\text{hat}\lambda\,(1+N)}{2\,\lambda}-\frac{N^2\,\lambda}{2\,\text{hat}\lambda\,(1+N)}}\sqrt{1+N}}{\sqrt{\text{hat}\lambda}\,\sqrt{2\,\pi}\,\lambda^{3/2}\,(-\text{Log}[\lambda\text{mn}]+\text{Log}[\lambda\text{mx}])}$$

(* The marginal distribution of the estimate with N=2500 *)

probhatλ

$$\frac{1}{2\,\text{hat}\lambda\,\text{Log}\left[\frac{\lambda\text{mx}}{\lambda\text{mn}}\right]}\left(-\text{Erf}\left[\frac{-2501\,\text{hat}\lambda+2500\,\lambda\text{mn}}{\sqrt{5002}\,\sqrt{\text{hat}\lambda\,\lambda\text{mn}}}\right]+e^{5000}\,\text{Erf}\left[\frac{2501\,\text{hat}\lambda+2500\,\lambda\text{mn}}{\sqrt{5002}\,\sqrt{\text{hat}\lambda\,\lambda\text{mn}}}\right]+\text{Erf}\left[\frac{-2501\,\text{hat}\lambda+2500\,\lambda\text{mx}}{\sqrt{5002}\,\sqrt{\text{hat}\lambda\,\lambda\text{mx}}}\right]-e^{5000}\,\text{Erf}\left[\frac{2501\,\text{hat}\lambda+2500\,\lambda\text{mx}}{\sqrt{5002}\,\sqrt{\text{hat}\lambda\,\lambda\text{mx}}}\right]\right)$$

(* This numerical integration takes too long and gives no hint of finishing in a reasonable time. We will investigate the comparison piecewise as motivated by plotting the extremes of the numerator/2 of probhatλ *)

(*numeratordiv2=
Numerator[

$$\frac{1}{2\,\text{hat}\lambda\,\text{Log}\left[\frac{\lambda\text{mx}}{\lambda\text{mn}}\right]}\left(-\text{Erf}\left[\frac{-2501\,\text{hat}\lambda+2500\,\lambda\text{mn}}{\sqrt{5002}\,\sqrt{\text{hat}\lambda\,\lambda\text{mn}}}\right]+e^{5000}\,\text{Erf}\left[\frac{2501\,\text{hat}\lambda+2500\,\lambda\text{mn}}{\sqrt{5002}\,\sqrt{\text{hat}\lambda\,\lambda\text{mn}}}\right]+\text{Erf}\left[\frac{-2501\,\text{hat}\lambda+2500\,\lambda\text{mx}}{\sqrt{5002}\,\sqrt{\text{hat}\lambda\,\lambda\text{mx}}}\right]-e^{5000}\,\text{Erf}\left[\frac{2501\,\text{hat}\lambda+2500\,\lambda\text{mx}}{\sqrt{5002}\,\sqrt{\text{hat}\lambda\,\lambda\text{mx}}}\right]\right)\Bigg]\Big/2$$ //.
{λmn→1,λmx→42643},
We want this term to equal one through its range,
which seems true except at the far ends of the range *)

```
Plot[numeratordiv2, {hatλ, 0.75, 1.25}, PlotTheme → "Scientific"]
```

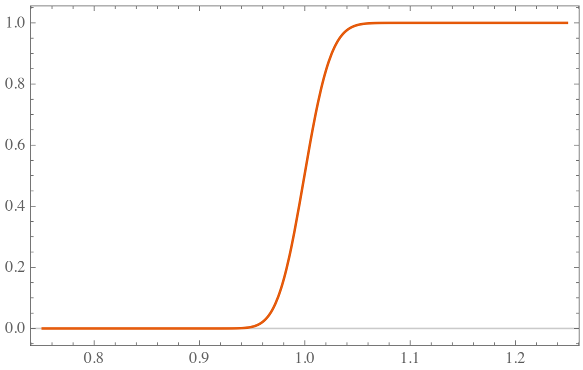

(* The range of interest is {λmn→1,λmx→42643} *)

```
Plot[numeratordiv2, {hatλ, 40 000, 42 643}, PlotTheme → "Scientific"]
```

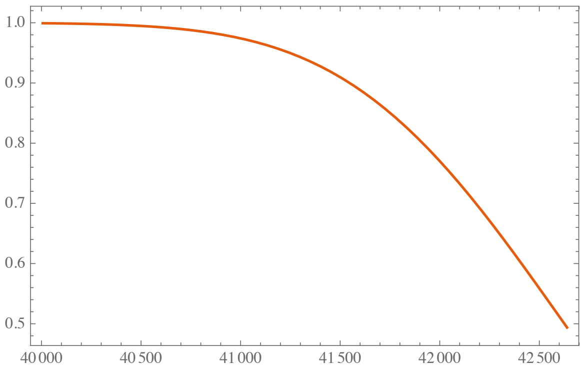

 William Levy, Victoria Calvert

```
(* This numerical integration takes too long and gives no
 hint of finishing in a reasonable time. We will investigate
 the comparison piecewise as motivated by plotting the
 extremes of the numerator/2 of probhatλ *)

(*numeratordiv2=
 Numerator[1/(2 hatλ Log[λmx/λmn])
   (-Erf[(-2501 hatλ+2500 λmn)/(Sqrt[5002] Sqrt[hatλ λmn])]+e^5000 Erf[(2501 hatλ+2500 λmn)/(Sqrt[5002] Sqrt[hatλ λmn])]+
    Erf[(-2501 hatλ+2500 λmx)/(Sqrt[5002] Sqrt[hatλ λmx])]-e^5000 Erf[(2501 hatλ+2500 λmx)/(Sqrt[5002] Sqrt[hatλ λmx])])]/2 //.
  {λmn→1,λmx→42643},
We want this term to equal one through its range,
which seems true except at the far ends of the range *)
```

In[805]:= `Plot[numeratordiv2, {hatλ, 0.75, 1.25}, PlotTheme → "Scientific"]`

Out[805]=

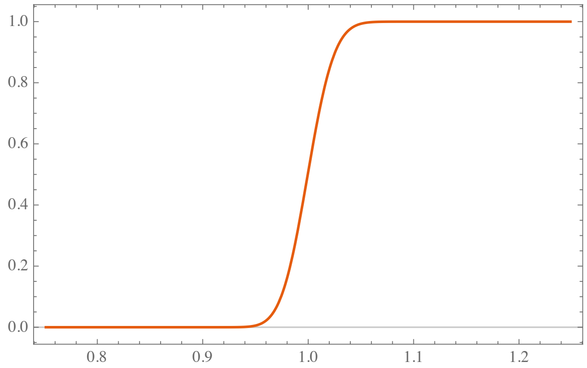

```
(*  The range of interest is {λmn→1,λmx→42643}   *)
```

In[806]:= `Plot[numeratordiv2, {hatλ, 40 000, 42 643}, PlotTheme → "Scientific"]`

Out[806]=

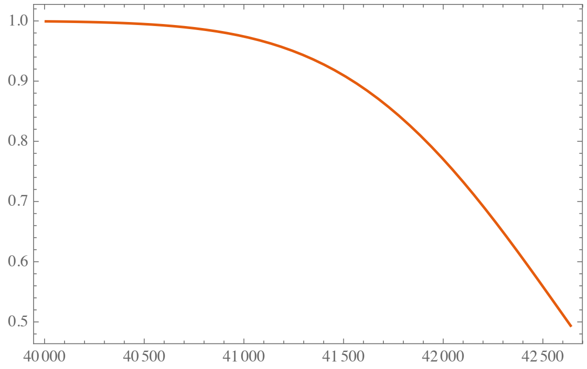

```
(* To evaluate the error we do numerical integration over
 the relevant subranges of each side of the densities *)

(* For the lefthand side *)

In[787]:= NIntegrate[probhatλ //. {λmn → 1, λmx → 42 643}, {hatλ, 1, 1.05}]

Out[787]= 0.00385112

In[788]:= NIntegrate[pλ //. {λmn → 1, λmx → 42 643}, {λ, 1, 1.05}]

Out[788]= 0.00457667

(* A difference of 7.2 parts in 10000 relative to the entire
 distribution *)

In[793]:= 0.004576672973368834` - 0.003851118693082693`

Out[793]= 0.000725554

(* And next show that there is little reason to consider a
 wider range because the difference is so small as the
 integration is moved to larger,
successively adjacent partitions *)

In[790]:= NIntegrate[probhatλ //. {λmn → 1, λmx → 42 643}, {hatλ, 1.05, 1.55}]

Out[790]= 0.0365288

In[791]:= NIntegrate[pλ //. {λmn → 1, λmx → 42 643}, {λ, 1.05, 1.55}]

Out[791]= 0.036533

In[794]:= 0.03653303698523965` - 0.03652878231245059`
```

Out[794]= $4.25467 \cdot 10^{-6}$

```
In[809]:= NIntegrate[probhatλ //. {λmn → 1, λmx → 42 643}, {hatλ, 1.55, 15.5}]

Out[809]= 0.21599

In[810]:= NIntegrate[pλ //. {λmn → 1, λmx → 42 643}, {λ, 1.55, 15.5}]

Out[810]= 0.21599

In[811]:= % - %%

Out[811]= 0.
```

```
(* Now examine the righthand side in the same manner *)

In[795]:= NIntegrate[probhatλ //. {λmn → 1, λmx → 42 643}, {hatλ, 40 000, 42 643}]

Out[795]= 0.00523492

In[796]:= NIntegrate[pλ //. {λmn → 1, λmx → 42 643}, {λ, 40 000, 42 643}]

Out[796]= 0.00600187

(* Again, a little over 7 parts in 10000 discrepancy *)

In[797]:= 0.00600187318068697` - 0.005234924845650944`

Out[797]= 0.000766948

(* and little further discrepancy accumulates if this righthand
 range is extended *)

In[798]:= NIntegrate[probhatλ //. {λmn → 1, λmx → 42 643}, {hatλ, 20 000, 40 000}]

Out[798]= 0.065019

In[799]:= NIntegrate[pλ //. {λmn → 1, λmx → 42 643}, {λ, 20 000, 40 000}]

Out[799]= 0.0650194

In[800]:= 0.06501941573344873` - 0.06501904134217998`

Out[800]= 3.74391 · 10^-7

(* To obtain the estiamted total absolute discrepancy between
 the distributions ,
add the left and right ones *)

In[807]:= 0.0007255542802861409` + 0.0007669483350360258`

Out[807]= 0.0014925

(* ca. 1.5 parts in  1000 *)
```

Energy Audit Overview Including Partitioning

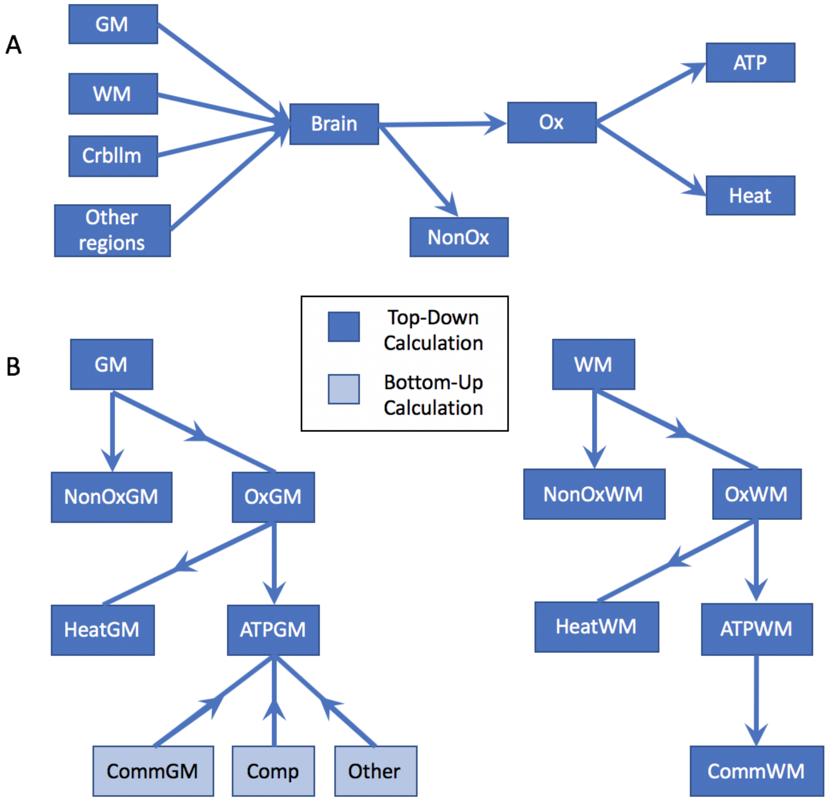


**Fig. S1.** The partitioning of the energy available from glucose for the top-down estimates. Note: Arrows indicate our partitioning process, not flow of energy/glucose. A: Proceeding from left to right, total brain glucose-uptake is calculated by summing the regional uptakes. From there, the total glucose is subdivided by metabolic fate. B: We begin with the regional cortical gray matter glucose uptake and end with specific ATP consumption. The first level of partitioning combines regional rates of [$^{11}$C]glucose uptake with regional brain weights; at the second level, glucose is partitioned based on metabolic fate (OxGM, the glucose that goes into cellular respiration, and NonOxGM, the glucose that is not oxidized); at the third level, energy from oxidized glucose is partitioned between ATP production and heat generation, and the majority of glucose-energy goes to the latter. At the fourth level, ATP energy is partitioned between communication, computation, and *Other*. GM-gray matter; WM-white matter; Crbllm-cerebellum; Other regions, see Table S1; NonOx-nonoxidized glucoxe; Ox-oxidized glucose; Comm-communication; Comp-computation; Other-includes synaptic modification, growth/retraction, maintenance, etc.

 

#### Table S1. Glucose Partitioning

| | Mass (g) ◇ | Glucose uptake ◇◇ $\mu mol/100g/min$ | Glucose uptake * $\mu mol$/region/sec |
|---|---|---|---|
| whole brain | 1495 | – | 6.05** |
| forebrain cortex | | | |
| cortical gray | 633 | 28.6 | 3.02 |
| cortical white | 590 | 18.4 | 1.81 |
| cerebellum | 154 | 24.7 | 0.63 |
| other regions † | 118 | 30†† | 0.59 |
| ventricular‡ | 5-15 | 0 | |

◇ Regional masses from (1).
◇◇ values from (2)
* Individual regional values are calculated from first two columns.
** This is a sum of the regional values. See text for details.
† Includes basal ganglia, thalamus, hypothalamus, etc.
†† Uses avg. glucose uptake value of striatum and thalamus of (2)
‡ This range is based on the possible remaining mass using values from (1)

#### Table S2. Glucose Energy Partitioning to ATP-watts◇

| Top-Down Calculations | Watts◇ (complete oxidation) | Non-oxidized◇◇ (equivalent watts) | ATP-watts* | ATP-watts** |
|---|---|---|---|---|
| whole brain (1495 g) | 17.0 | 1.86 | 6.19 | 5.23 |
| cerebellum (154 g) | 1.77 | 0.19 | 0.65 | 0.55 |
| other regions (118 g)† | 1.65 | 0.18 | 0.60 | 0.51 |
| forebrain cortex (1223 g): | | | | |
| white (590 g) | 5.07 | 0.56 | 1.85 | 1.56 |
| gray (633 g) | 8.45 | 0.93 | 3.09 | 2.61 |
| Bottom-up calculations | | | | |
| gray: 1.6Hz | | | | |
| communication | | | 2.75 | 2.75†† |
| computation | | | 0.17 | 0.17 |
| other ATP demands | | | 0.17‡ | 0.17‡ |
| gray: 2.5Hz | | | | |
| communication | | | 3.29†† | 3.29†† |
| computation | | | 0.25 | 0.25 |
| other ATP demands | | | 0.25‡ | 0.25‡ |

◇ Watts based on glucose-uptake values from (2) and 2.8 $MJ/mol\ glucose$ (23) and 36 $kJ/molATP$ (5)(6); regional masses from (1).
◇◇ Also assuming complete oxidation of glucose. See partitioning of glucose in earlier sections
* Using Nath's torsional mechanism (5), (24) which incorporates mitochondrial leak
** Using chemiosmotic theory (25) which is then downgraded by standard mitochondrial leak value: 20% (26)
† Including basal ganglia, thalamus, brainstem, etc. The missing mass is ventricular. See Table S1 and the accompanying footnotes for more information.
†† Indicates bottom-up values exceed available energy if the top-down calculations are accepted.
‡ Assuming that "other" consumes the same amount of energy as computation.

**Table S3. Bottom-Up Computation and Communication**$^{\star}$

| Gray Matter Computational Costs$^{\diamond}$ | | |
|---|---|---|
| AMPAR | NMDAR$^{\dagger}$ | AMPAR + NMDAR |
| 0.113 W | $0.5 \cdot 0.113 \approx 0.057$ W | 0.17 W |
| **Gray Matter Communication Costs** | | |
| Resting Potential | Action Potentials | Presynaptic Transmission |
| 1.81 W | 0.75 W | 0.19 W |
| $7.5 \cdot 10^{-11}$ J/NRN/AP | $3.1 \cdot 10^{-11}$ J/NRN/AP | $7.9 \cdot 10^{-12}$ J/NRN/AP |
| **Gray Matter Communication + Computation** | | |
| GM Comm Total | Comp Total | Comm+Comp |
| 2.75 W | 0.17 W | 2.92 W |
| $1.14 \cdot 10^{-10}$ J/NRN/AP | $7.1 \cdot 10^{-12}$ J/NRN/AP | $1.94 \cdot 10^{-10}$ W/NRN |
| **Other Totals (cortex = WM + GM)** | | |
| WM$^{\star\star}$+ GM Comm | Comp + *Other*$^{\diamond\diamond}$ | Total GM + WM |
| 4.6 W/cortex | 0.34 W | 4.94 W/cortex |
| $1.92 \cdot 10^{-10}$ J/NRN/AP | $1.4 \cdot 10^{-11}$ J/NRN/AP | $2.1 \cdot 10^{-10}$ J/NRN/AP |

$^{\star}$ For more details on these calculations, see Methods
$^{\diamond}$ 1.6 Hz with 75% failure rate implies 0.4 SA/synapse/sec
$^{\dagger}$ Assumes NMDA-receptor activation contributes half-again the cost of AMPA-receptor activations
$^{\star\star}$ White matter communications also includes white matter *Other*
$^{\diamond\diamond}$ Gray matter *Other* includes synaptic modification costs, e.g., metabotropic transmitter effects, axonal growth/retraction, receptor modification/removal as well as the transport needed for such modifications

**Table S4. Computational costs arising from ionotropic glutamate synaptic activations**

| Voltages | | | |
|---|---|---|---|
| $V_{rev}$<br>$-7$ mV | $V_{m,ave}$<br>$-55$ mV | $V_{Na,Nern.}$<br>$+55$ mV | $V_{K,Nern}$<br>$-90$ mV |
| For a single synaspe, Average (Ave) AMPA-Receptor (AR) per synaptic activation (SA)$^{\star}$ | | | |
| $G_{ave}$<br>200 pS/SA | $(V_{Na^+} - V_{ave,m}) \cdot G_{Na}$/SA<br>110 mV $\cdot 114.5$ pS/SA | $Na^+$ amps<br>12.5 pA/SA | Na$^+$ coulombs/SA<br>15.1 fC/SA |
| AMPAR-based (All GM synapses$= 1.5 \cdot 10^{14}$) | | | |
| $Na^+$flux<br>3.6 C/sec | $Na^+$flux<br>$3.8 \cdot 10^{-5}$ mol/sec | ATP mol/sec<br>$1.3 \cdot 10^{-5}$ mol/sec | ATP watts<br>0.113 W |
| AMPAR + NMDAR Computational cost | | | |
| per cerebral cortex<br>$1.5^{\dagger} \cdot 0.113$ W $\approx 0.17$ W | | per neuron per spike<br>$7.1 \cdot 10^{-12}$ J | |

$^{\star}$ SA duration 1.2 msec
$^{\diamond}$ 1.6 Hz with 75% failure rate implies 0.4 SA/synapse/sec
$^{\dagger}$ Assumes NMDA-receptor activation contributes half-again the cost of AMPA-receptor activations

#### Table S5. Gray Matter Communication$^{\diamond}$

| Parameters. I (all conductances are at rest) | | | |
|---|---|---|---|
| $V_m^{rest}$ | $V_{Na,Nern}$ | $V_{K,Nern}$ | $G_{Na} : G_K$ |
| $-66\, mV$ | $+55\, mV$ | $-90\, mV$ | $121 : 24$ |
| Parameters. II | | | |
| Capacitivity | Resistivity | Conductivity | Axon/Bouton Area |
| $0.96\, \mu F/cm^2$ | $30,000\, \Omega cm^2$ | $3.33 \cdot 10^{-5}\ S/cm^2$ | 2:1 |
| Total across all axons or boutons | | | |
| Area Axon + Bouton | $G^{rest}_{axon+btn}$ | $G^{rest}_{Na}$ | $C_{ax}; \quad C_{btn}$ |
| $21.8 \cdot 10^6\ cm^2$ | $727\, S$ | $111\, S$ | $14.6\, F;\ 6.34\, F$ |
| $Na^+$ rest-flux and ATP to remove | | | |
| $Na^+$ flux | $Na^+$ flux | ATPs used | |
| $14.6\, C/s$ | $1.51 \cdot 10^{-4}\ mol/s$ | $5.03 \cdot 10^{-5}\ mol/s$ | |
| Cost of axon + bouton rest potentials ($36000\ J/mol ATP$ ) | | | 1.81W |
| 110mV ($V_{AP}$) Axon Action Potential (AP); $20\ mV$ ($\Delta V_{Bt}$) Bouton depolarization | | | |
| Axon charging/sec$^{\diamond}$ | Overlap-scaled$^{\star}$ | Bouton charging/sec$^{\diamond\diamond}$ | $Na^+ mol/sec$ |
| $2.57$ amps | $5.86$ amps | $0.20$ amps | $6.3 \cdot 10^{-5}$ mol/sec |
| Cost of action potentials ($2.09 \cdot 10^{-5}\ ATP\, mol/sec$) | | | 0.75W |
| ATP per vesicle released (VR); ATP per $Ca^{2+}$ spike | | | |
| $ATP/vesicle$ | $Ca^{+2}\ removal$ | | |
| $5.7 \cdot 10^3$ ATPs | $1.2 \cdot 10^4$ATPs | | |
| ($9.5 \cdot 10^{-21}$ mols) | ($1.99 \cdot 10^{-20}$ mols) | | |
| Presynaptic AP costs $^{\star\star}$ | | | |
| VR Events/s | Ca events/s | VR ATP use | $Ca^{2+}$ ATP use |
| $6.0 \cdot 10^{13}$/s | $2.4 \cdot 10^{14}/s$ | $5.7 \cdot 10^{-7}$mol/s | $4.78 \cdot 10^{-6}$ mol/s |
| AP Generated Presynaptic Cost | | | 0.19 W |
| Total | | | 2.75 W |

$^{\diamond}$ 1.6 Hz firing rate
$^{\star}$ Multiplier from (27)
$^{\star\star}$ 75% failure rate of TR but no failure of Ca entry

**Table S6. GM Communication Volume Fractions and Cylindrical Approximations**$^{\diamond}$

| | Vol. Frac | Volume $^{\diamond\diamond}$ | Diameter | Height* per bouton | Length** (total) | $Area_{pm}$ $^{\dagger}$ | $C_m$ $^{\dagger\dagger}$ |
|---|---|---|---|---|---|---|---|
| Boutons | 22% | 139 $cm^3$ | 1.1 $\mu$m | 1.0 $\mu$m | - | $6.61{\cdot}10^6$ $cm^2$ | 6.34 F |
| Axons | 30% | 190 $cm^3$ | 0.50 $\mu$m | - | $9.66{\cdot}10^{10}$ cm | $15.2 \cdot 10^6$ $cm^2$ | 14.6 F |
| Total | 52% | 329 $cm^3$ | - | - | - | $21.8 \cdot 10^6$ $cm^2$ | 20.9 F |

$^{\diamond}$ Assuming $1.5 \cdot 10^{14}$ synapses per cortex
$^{\diamond\diamond}$ Cortical gray matter volume
* Height is for a single cylindrical bouton
** Length is total length of all small axons
$^{\dagger}$ Area of plasma membrane (pm)
$^{\dagger\dagger}$ Membrane capacitance using 0.96 $\mu F/cm^2$ based on 0.88 $\mu F/cm^2$ of the lipid membrane plus activation of two-thirds of the 0.12 $\mu F/cm^2$ of $Na^+$channel gating-charge.

**Table S7. Other axon parameter sets consistent with a 30% volume-fraction**

| Axon Diameter ($\mu m$) | 0.28 | 0.4 | 0.5 | 0.6 |
|---|---|---|---|---|
| Axon Area ($cm^2$) | $27.1 \cdot 10^6$ | $18.9 \cdot 10^6$ | $15.2 \cdot 10^6$ | $12.7 \cdot 10^6$ |
| Total Area$^{\diamond}$($cm^2$) | $33.7 \cdot 10^6$ | $25.6 \cdot 10^6$ | $21.8 \cdot 10^6$ | $19.3 \cdot 10^6$ |
| Axon Length$^{\diamond\diamond}$($cm$) | $3.08 \cdot 10^{11}$ | $1.51 \cdot 10^{11}$ | $9.67 \cdot 10^{10}$ | $6.72 \cdot 10^{10}$ |
| RP cost (W) | 2.79 | 2.13 | 1.81 | 1.60 |
| AP cost (W) | 1.32 | 0.93 | 0.75 | 0.63 |

$^{\diamond}$ Total area is the sum of axonal surface area and bouton surface area as in Table S6, 22% volume-fraction.
$^{\diamond\diamond}$ The axon length is the length of all axons extended by 1.0 $\mu$m assumed bouton height and number.

**Table S8. Effect of Varying Bouton Dimensions**

| Diameter | Height | $Area_{pm}$ | Vol. Frac | $C_m$ |
|---|---|---|---|---|
| 0.9$\mu m$ | 1.0$\mu m$ | $5.19 \cdot 10^6 cm^2$ | 15% | $4.99F$ |
| 0.9$\mu m$ | 1.2$\mu m$ | $6.04 \cdot 10^6 cm^2$ | 18% | $5.80F$ |
| $1.1\mu m$ | 1.0$\mu m$ | $6.61 \cdot 10^6 cm^2$ | 22% | $6.34F$ |
